# CHI3L1-Targeted Small Molecules as Glioblastoma Therapies: Virtual Screening-Based Discovery, Biophysical Validation, Pharmacokinetic Profiling, and Evaluation in Glioblastoma Spheroids

**DOI:** 10.1101/2025.05.13.653859

**Authors:** Hossam Nada, Longfei Zhang, Baljit Kaur, Moustafa T. Gabr

**Affiliations:** Department of Radiology, Molecular Imaging Innovations Institute (MI3), Weill Cornell Medicine, New York, NY 10065, USA

**Author notes:** To whom correspondence should be addressed: Moustafa T. Gabr.

**Keywords:** Virtual screening, Biophyiscal validation, Glioblastoma, Pharmacokinetics, 3D spheroids

## Abstract

Glioblastoma (GBM) remains the most aggressive primary brain malignancy with a 10% three-year survival rate. Chitinase-3-like protein 1 (CHI3L1) has emerged as a critical factor in the progression of GBM progression, invasion, and treatment resistance. However, small molecule inhibitors targeting CHI3L1 are largely unexplored. Microscale thermophoresis (MST) investigation of the direct binding potential of reported CHI3L1 modulators (**K284, G721-0282, CHI3L1-IN-1**) revealed modest to undetectable direct CHI3L1 binding affinity. Herein, pharmacophore-based virtual screening of in-house library resulted in the discovery of **G28** as the most potent small molecule CHI3L1 binder reported to date. The CHI3L1 binding affinity of **G28** was validated using MST and surface plasmon resonance (SPR). To evaluate the GBM-modulatory potential of **G28**, we conducted comprehensive pharmacokinetic and 3D spheroid studies alongside established CHI3L1 modulators. **G28** demonstrated outstanding bioavailability and low toxicity, addressing key limitations faced by previous CHI3L1-targeted strategies. Notably, in 3D GBM spheroid models, **G28** significantly outperformed reported CHI3L1 small molecule modulators, showing the most pronounced dose-dependent reductions in spheroid weight, migration, and viability. These findings position **G28** as the most promising CHI3L1-targeted small molecule to date and a compelling candidate for GBM therapeutic development.

---

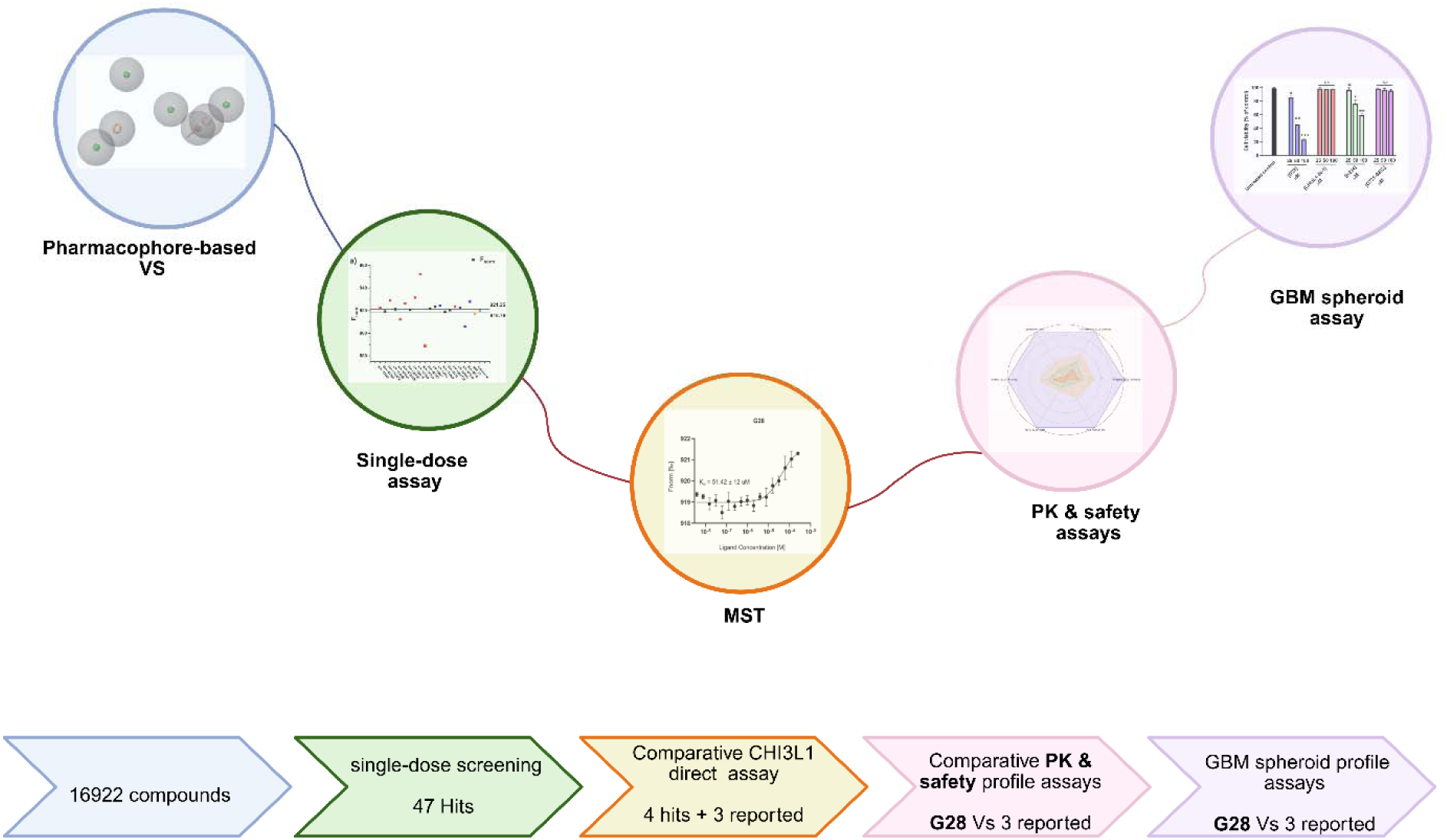

## 1. Introduction

Among brain tumors, glioblastoma (GBM) is widely recognized as the most aggressive and lethal primary brain malignancy which is characterized by a with a 3-year survival rate of only around 10%^1^.

The median survival time remains short due to the aggressive nature of the GBM and its resistance to traditional treatments as well as the immunosuppressive microenvironment^2^. The current standard treatment for GBM includes maximal surgical resection followed by radiation therapy and temozolomide(TMZ)^3, 4^ chemotherapy which is often combined with tumor-treating fields. Surgical resection aims to remove as much tumor tissue as possible, but microscopic infiltrative cells often persist which often leads to recurrence^5, 6^.

Despite advances, GBM therapy is hindered by multiple factors such as the limited systemic drug penetration by the BBB where only about 20% of systemically administered temozolomide reaches the cerebral parenchyma^7, 8^. The efflux pumps like ABC transporters^9, 10^ further reduce intracellular drug concentrations. Moreover, the recurrence of GBM is highly recurrent due to the presence of glioma stem cells (GSCs), which possess high migratory potential, resistance to chemotherapy and radiation in addition to the ability to form recurrent tumors^11, 12^. Immunotherapies struggle in GBM’s "cold tumor" microenvironment, characterized by low T-cell infiltration and high levels of immunosuppressive cytokines like TGF-β. Even with aggressive treatments, median survival remains 12–18 months, underscoring the need for urgent and critical need to develop novel therapeutic strategies for GBM that can overcome the limitations of existing treatment modalities.

Among the molecular targets implicated in glioblastoma (GBM), chitinase-3-like protein 1 (CHI3L1, also known as YKL-40) has emerged as a key factor in GBM pathogenesis^13, 14^. CHI3L1 is a secreted glycoprotein that lacks enzymatic chitinase activity but still plays a fundamental role in promoting tumor progression by enhancing cell survival, migration, invasion, and angiogenesis^14–18^. In the brain, CHI3L1 is primarily expressed by astrocytes and microglia under the transcriptional control of regulators such as NFI-X3 and STAT3^19^. CHI3L1 exerts its effects through interactions with a network of receptors including CRTH2^20^, RAGE^21^, IL-13 receptor α2 (IL-13Rα2)^22^, PAR-2^23^, and CD44^24^. The interaction of CHI3L1 with its receptors triggers the activation of key oncogenic signaling pathways—including ERK1/2, PI3K/Akt, Wnt/β-catenin, JNK, and NF-κB—which collectively govern critical processes such as inhibition of apoptosis, modulation of immune responses, extracellular matrix remodeling, and inflammasome activation^25, 26^.

Studies have shown that CHI3L1 is highly expressed in glioblastoma^27^ compared to other cancers and normal tissues. Additionally, studies haves shown that the expression of CHI3L1 correlates with clinical and molecular features of malignancy where high CHI3L1 expression have been associated with poor patient prognosis, resistance to chemotherapy and radiotherapy^28^. Additionally, Chi3L1 expression levels have been established as a key marker for the mesenchymal subtype of glioblastoma^29^. At the molecular level, CHI3L1 has been found to regulate glioma cell invasion, migration, growth, and tumor vascularization^30^. Furthermore, recent studies have indicated that CHI3L1 plays important roles in the glioblastoma immune microenvironment where it closely associated with immune responses, inflammatory activities, and immunosuppression^1, 31^. Meanwhile, inhibition of CHI3L1 have been reported to reduce immunosuppression, decrease the mesenchymal signature of glioblastoma, and restrict GSC plasticity, potentially overcoming immunotherapy resistance^32^. Together, these findings suggest that CHI3L1-targeted immunotherapy, either alone or in combination with other immunotherapies such as immune checkpoint inhibitors, represents a promising new strategy for treating GBM by modulating cellular plasticity and reducing tumor burden.

To date, only a few small molecules CHI3L1 modulators have been reported; **K284**^33^, **G721- 0282**^34^ (**G721**), and **CHI3L1-IN-1**^35^ (Figure 1). CHI3L1-IN-1 was identified via an indirect AlphaScreen assay, which is effective for identifying compounds that compete with a known probe but does not confirm direct binding or reveal binding modes. Meanwhile, K284 was discovered through pull-down assays and G721 was proposed based on docking studies, important gaps remain in our understanding of their mechanisms of action and their capacity to modulate CHI3L1 function effectively. Furthermore, assessment of the direct binding of these modulators using microscale thermophoresis (MST) resulted in a varied activity profile with **G721** exhibiting no detectable dose-dependent binding. Meanwhile, **CHI3L1-IN-1** exhibited weak binding with a dissociation constant (KD) in the millimolar range. Lastly, **K284** demonstrated measurable binding with a KD of 152 µM.

**Figure 1.**
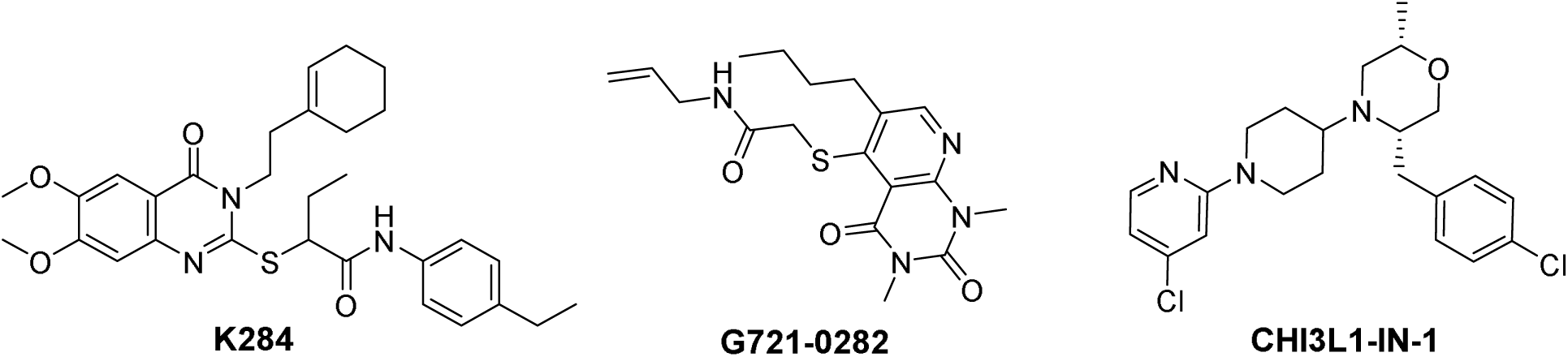
Chemical structure of the reported CHI3L1 inhibitors; **K284**, **G721-0282** (**G721**), and **CHI3L1-IN- 1**.

Given the weak to moderate activity of current CHI3L1 inhibitors, the discovery of more potent CHI3L1 modulators represents a critical unmet need. While traditional drug discovery pathways are resource-intensive, virtual screening (VS) offers an efficient alternative for early-stage hit identification^36,37^. Moreover, with recent crystallization of CHI3L1^35^ reported the presence of a distinct binding pocket, offering an opportunity to apply structure-based drug discovery approaches to identify more potent and selective modulators. Accordingly, we planned to carry out a VS assay to identify novel small molecules capable of modulating CHI3L1 with improved potency in comparison to current modulators. Moreover, given that ideal therapeutic candidates for GBM must combine biochemical potency with blood-brain barrier (BBB) penetration and favorable pharmacokinetic (PK) properties we planned for a PK assay for the top identified molecules.

Herein, we present a CHI3L1-targeted drug discovery pipeline that integrates pharmacophore-based virtual screening (VS) of an in-house small-molecule library composed of 16,922 small molecules to identify novel modulators of CHI3L1. The identified candidates were subjected to a side-by-side evaluation of CHI3L1-binding potency along with previously reported modulators using microscale thermophoresis (MST). The most promising hit from the VS campaign underwent detailed pharmacokinetic (PK) profiling and was further evaluated for efficacy in GBM using three-dimensional (3D) spheroid models.

3D spheroid models were employed for their enhanced physiological relevance which offers a more accurate representation of the tumor microenvironment (TME) compared to traditional two-dimensional (2D) monolayer cultures^38^. The increased accuracy of 3D spheroid models is due to the fact that 3D cultures allow cancer cells to organize in spatially relevant architectures which provides a good representation of the structure and signaling dynamics of native tumors^39^. Conversely, in 2D cell models the cells are grown as flat monolayers leading to poor representation of critical cell–cell and cell–matrix interactions. Cells in 2D culture often undergo cytoskeletal rearrangements and acquire artificial polarity, leading to aberrant gene and protein expression. In contrast, 3D-cultured cells exhibit more physiologically relevant morphological and molecular characteristics, making them a superior platform for evaluating therapeutic efficacy in vitro^40–42^. The PK and GBM spheroid models were extended to all known CHI3L1 modulators, establishing the first standardized comparison of their binding affinities and functional effects in disease-relevant systems. This integrated approach establishes a robust platform for rational CHI3L1 modulators discovery with high translational potential.

## 2. Results and Discussion

### 2.1. Virtual screening

Virtual screening of the two in-house libraries was carried out in two sequential steps. The first involved pharmacophore screening followed by ranking of the resulting hits using a three-step docking based protocol. In order to carry out the pharmacophore screening, two distinct pharmacophore hypotheses were generated. The first hypothesis composed of a seven-feature ligand-based hypothesis derived from compound **CHI3L1-IN-1** (Figure 2a), a recently reported CHI3L1 inhibitor. While the second pharmacophore exhibited a ten-feature structure-based hypothesis generated from the CHI3L1 binding site.

**Figure 2.**
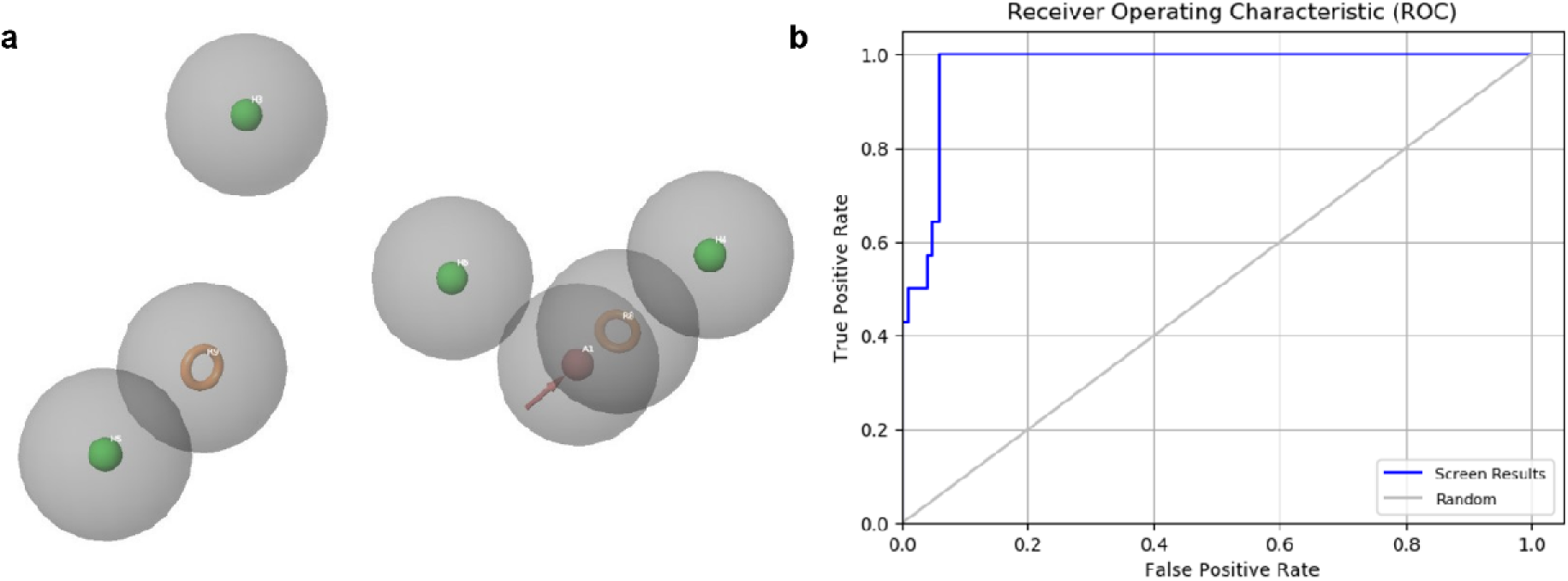
Ligand-based pharmacophore model used for virtual screening. (**a**) E-pharmacophore hypothesis with key pharmacophoric features (AHHHHRR). (**b**) Receiver operating characteristic (ROC) curve analysis evaluating the predictive performance of the ligand-based pharmacophore model.

Due to the limited availability of known CHI3L1 inhibitors, 100 decoys were generated via the DUDE server to evaluate both models. Meanwhile, known actives from the same study as compound X were used for validation. Evaluation of the structure-based hypothesis showed its inability to identify any active compounds even when the similarity threshold was relaxed to four features to improve sensitivity. In contrast, the ligand-based hypothesis exhibited promising predictive power when the screening was subjected to a criterion of matching at least five of seven pharmacophore features. The ligand-based hypothesis consisted of seven features consisting of two aromatic rings (R9 and 9), four hydrophobic regions (H3-6) and a H-bond acceptor site (A1).

Evaluation of the ligand-based hypothesis yielded a receiver operating characteristic (ROC) value of 0.97 (Figures 2b) and area under the accumulated curve (AUAC) score of 0.91. However, this threshold introduced false positives leading to the misclassification of few decoys as actives. In order to avoid any false positives in the VS, the ligand-based model was subjected to a stricter six-out-of-seven feature match which prioritized increased specificity. While upon reevaluation of the model while applying the stricter screening criteria led to the reduction of the ROC value to 0.67 which represents a moderate decrease in overall performance metrics. Notably, however, no false positives (decoys) were identified under these more stringent conditions. This outcome suggests an improvement in the model’s specificity, which is particularly valuable in the context of virtual screening campaigns where the prioritization of genuine active compounds is essential. While the decreased ROC value indicates some reduction in sensitivity, the elimination of false positives demonstrates an enhanced ability to discriminate between active compounds and decoys. As such, the trade-off between sensitivity and specificity was deemed to be advantageous for our screening. Consequently, the six-out-of-seven pharmacophoric match criterion was selected for screening the 16,922 in-house compounds to balance sensitivity with specificity, maximizing accuracy while minimizing false positives.

Compounds identified from the pharmacophore screening which matched at least six out of the seven defined pharmacophoric features were ranked using Maestro Schrödinger’s High-Throughput Virtual Screening (HTVS) module. In the HTVS, all pharmacophore-matched compounds were docked and ranked based on their predicted binding to the prepared crystal structure of CHI3L1 (PDB ID: 8R4X). The HTVS involved a three-step scoring and elimination iteration where compounds are progressively filtered based on their docking scores and binding affinity. A combination of HTVS results and visual inspection led to the selection of 47 compounds, sourced from two in-house libraries: Enamine (Table S1) and MedChemExpress (Table S2). These hits were then subjected to biophysical screening to validate the results.

### 2.2. Biophysical evaluation of CHI3L1 binding affinity

#### 2.2.1. Single dose screening

To evaluate the binding abilities of 47 selected hits from the virtual screening assay, single-concentration TRIC assays were conducted in singlet at 250 μM. Compounds exhibiting F_norm_ values outside the negative control range (defined as reference ± 3 × standard deviation) were retained for further analysis. The known CHI3L1 binder **CHI3L1-IN-1** was selected as a positive control to validate assay performance. Based on F_norm_ values, fifteen potential hits were identified which included 12 hits from the MedChemExpress library and 3 hits from the Enamine library (Figure 3), indicated in blue and red). Subsequently, TRIC-based autofluorescence and quenching assays were performed to eliminate candidates exhibiting signal interference or fluorophore interactions. This led to the identification of seven compounds; HY-15590, HY-108347, HY-103696, HY-19914 (hereafter referred to as **G28**), **E4**, **E11**, and **E15** (Figure 3, blue spots). The seven compounds showed no significant differences in initial fluorescence compared to the negative control (Figure S1).

**Figure 3.**
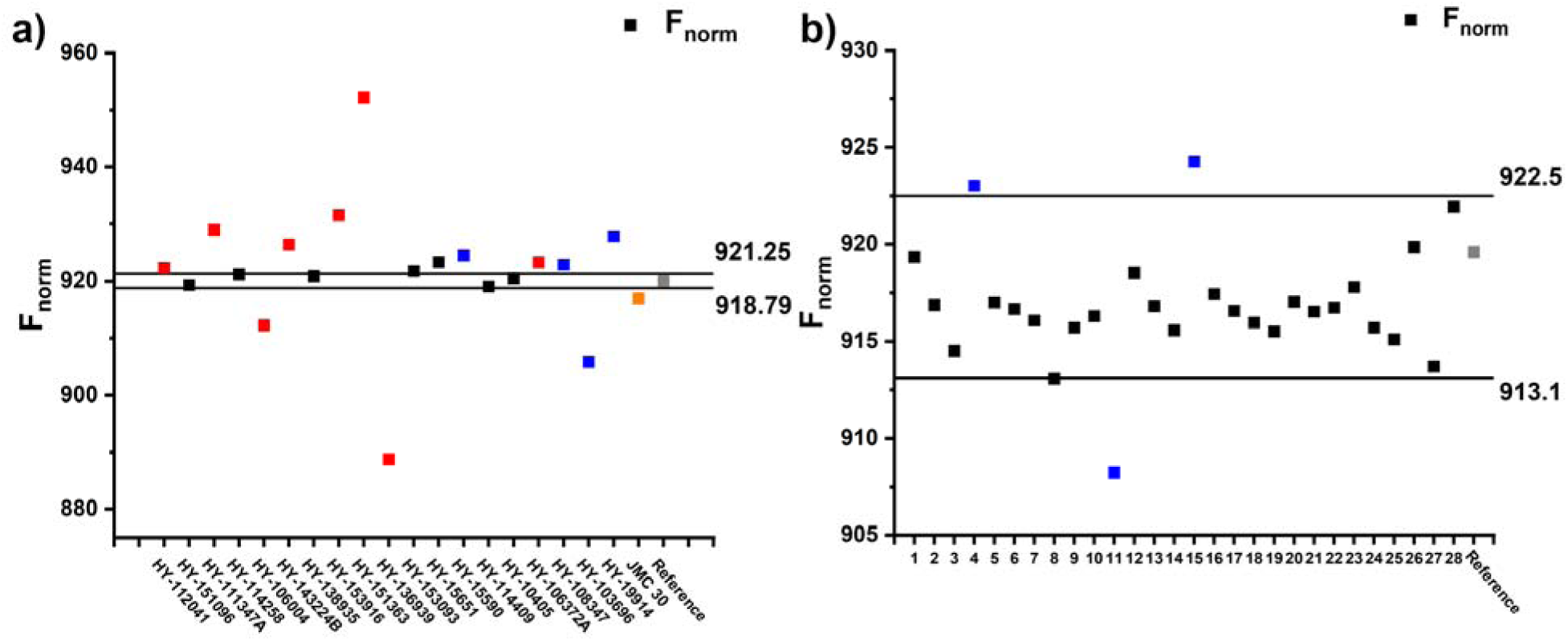
TRIC-based affinity screening of virtual candidates. Single dosage screening of molecules from the MedChemExpress library (**a**) and Enamine library (**b**). 250 μM (with 2.5% DMSO) compound was incubated with His-labeled hCHI3L1 to study the binding ability towards this protein. The blue, red, black, orange, and gray indicates the potential hits, molecules with autofluorescence or dye-interaction, non-binder, positive control (**J30**), and negative control (buffer with 2.5% DMSO). The black lines indicate the value of Mean_negative control_ ± 3 × SD. F_norm_ values were measured in triplicate with results given as the mean value. Buffer condition: 10 mM HEPES, 150 mM NaCl, 1% Pluronic F127, 1 mM TCEP, pH = 7.4. Incubation time: 30 min.

#### 2.2.2. Dose-dependent assay

The seven potential hits from the single-dose assay, along with **K284**, **G721** and **CHI3L1-IN-1** were evaluated in a dose-dependent assay to assess their direct binding potential with CHI3L1. Four of the seven potential hits (**G28**, **E4**, **E11**, and **E15**) exhibited dose dependent response which indicates that these are true hits rather than artifacts (Figure 4). The 2D structure of the four hits is illustrated in Figure S2. Among the investigated compounds, **G28** demonstrates the strongest binding with a KD of 51.42 μM, followed by **E4** (138 μM), **K284** (152 μM), **E15** (292 μM), and **E11** (390 μM), with **CHI3L1-IN-1** showing remarkably weaker binding in the millimolar range (77.3 mM). Meanwhile, **G721** failed to show any dose dependent binding when evaluated against CHI3L1 (Figure S3). The results indicate that **G28** is a promising lead for modulating CHI3L1 with improved binding profile than any previously reported CHI3L1 binders.

**Figure 4.**
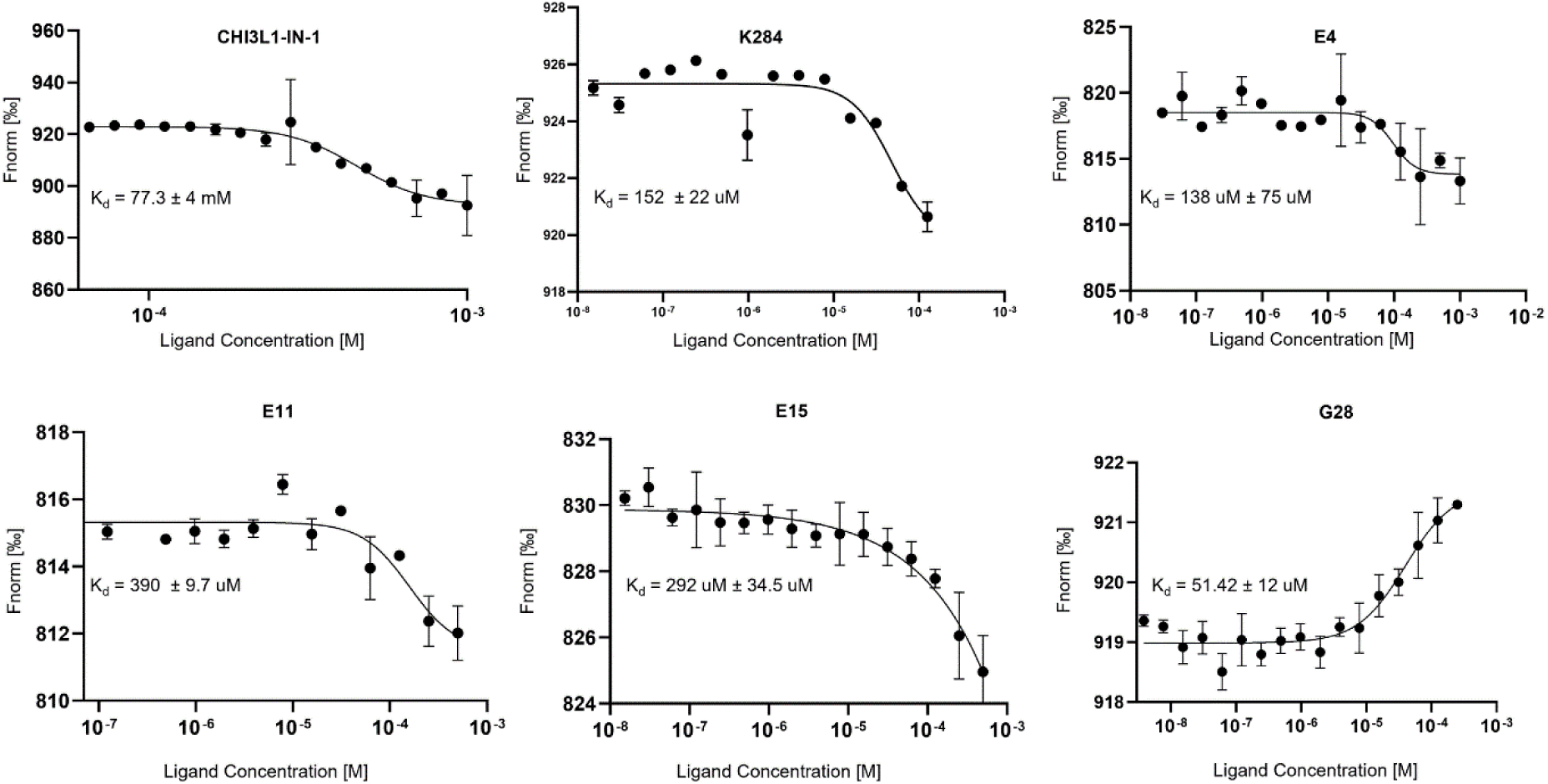
Direct binding validation of small molecule hits to CHI3L1 using MST. Dose-dependent binding curves for newly identified hits (**G28**, **E4**, **E11**, and **E15**) in comparison to previously reported modulators (**CHI3L1-IN-1** and **K284**) measured by MST. The normalized fluorescence (Fnorm [%]) is plotted against increasing ligand concentrations, with calculated dissociation constants (KD) indicated for each compound. **G28** demonstrates the strongest binding affinity (KD = 51.42 ± 0.24 μM), exhibiting a unique upward response curve compared to other compounds. **CHI3L1-IN-1** shows substantially weaker binding in the millimolar range (KD = 77.3 ± 4 mM). Data are presented as mean ± SEM from three independent experiments (n = 3) for all compounds except **K284** (n = 2).

### 2.3. SPR screening

To further validate the interaction between **G28** and CHI3L1, we performed surface plasmon resonance (SPR) as an orthogonal technique. SPR is well-established for its high sensitivity, ability to monitor real-time binding events, and label-free platform, rendering it an ideal complement to MST for confirming molecular interactions. In our SPR experiments, CHI3L1 was immobilized on the SPR chip and increasing concentrations of **G28** were flowed over the CHI3L1 surface. The resulting sensorgram (Figure 5) displayed a clear, dose-dependent increase in binding response, indicating specific interaction between **G28** and CHI3L1. This orthogonal validation by SPR not only corroborates the MST results but also strengthens the evidence that **G28** can engage CHI3L1.

**Figure 5.**
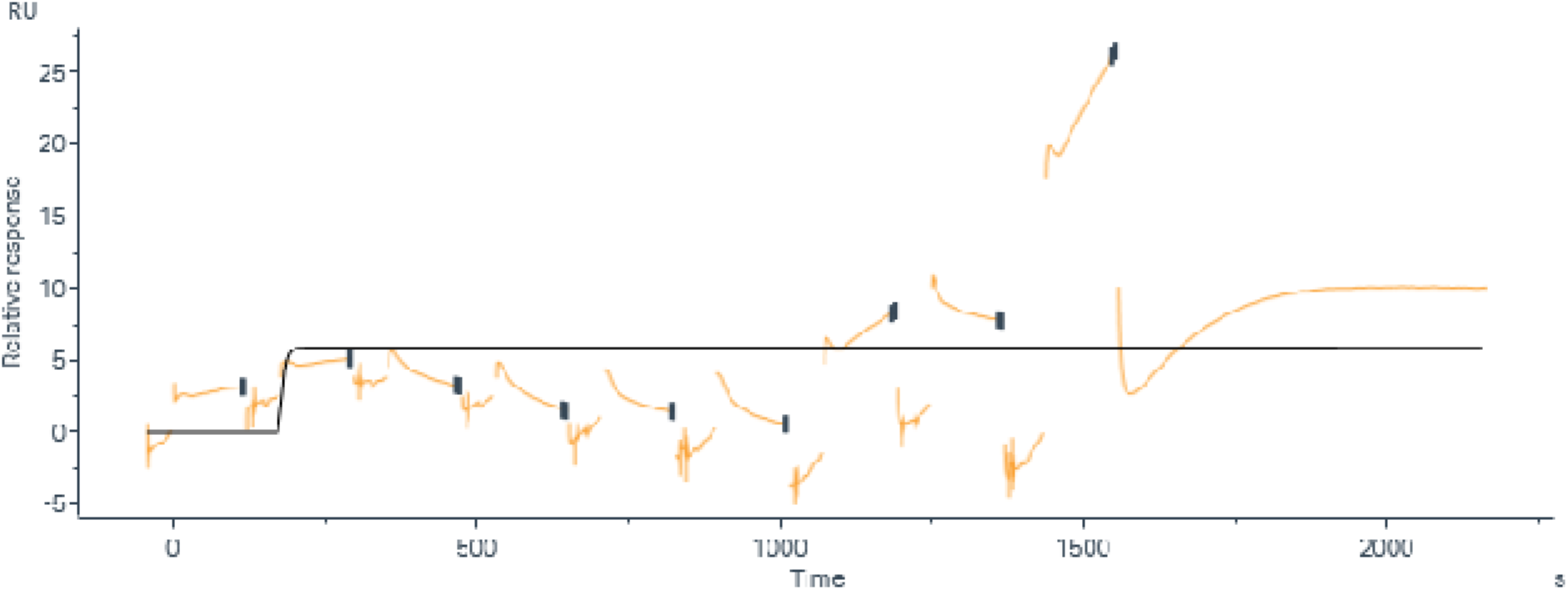
SPR validation of the CHI3L1 binding affinity of G28. Single-cycle kinetic sensorgram of **G28** interacting with the CHI3L1 protein using SPR.

### 2.4. In silico investigation

Molecular docking was conducted to investigate the difference between the binding modes of the top hit (**G28**) and the reported CHI3L1 binders (**K284, G721** and **CHI3L1-IN-1**). The four compounds illustrated a shared binding mode with CHI3L1 where they interacted with the same binding pocket (Figure 6A) indicated a shared binding mechanism despite their structural differences. The binding site appears to be a well-defined cavity that accommodates these structurally diverse molecules through various non-covalent interactions. The molecular docking investigation showed that **CHI3L1-IN-1** (depicted in green, Figures 6B-C) in complex with CHI3L1 displayed a network of favorable interaction with both hydrophobic and hydrophilic residues. However, **CHI3L1-IN-1** failed to exhibit any hydrogen bonds with CHI3L1 which might explain the observed lowered binding activity when compared to **G28**.

**Figure 6.**
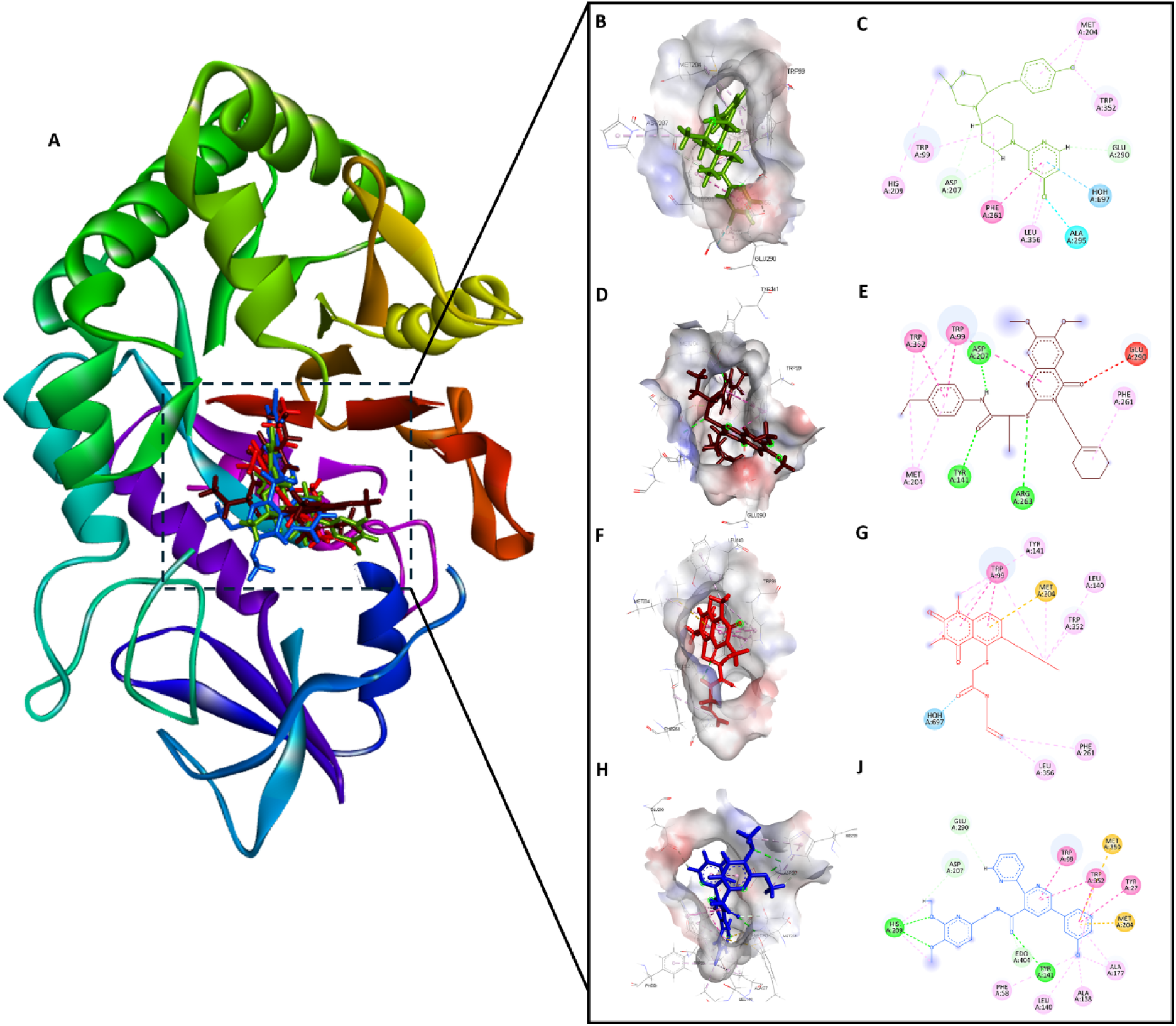
Molecular Docking of CHI3L1 with Inhibitors CHI3L1-IN-1 (green), **K284** (brown), **G721** (red) and **G28** (blue). (A) Superimposed view of the four compounds docked within the same binding site of CHI3L1. (B, D, F, H) 3D visualizations of the CHI3L1 in complex with **CHI3L1-IN-1**, **K284**, **G-721** and **G28**, respectively. (C, E, G, J) Corresponding 2D interaction diagrams between CHI3L1 residues and each compound. Key interactions are color-coded: green (hydrogen bonds), yellow (π–sulfur), orange (π– charge), dark pink (π–π stacking), purple (π–sigma), light pink (hydrophobic), red (unfavorable interactions) and light green (carbon-hydrogen bonds).

Meanwhile, **K284** (shown in brown, Figures 6D-E) was predicted to establish several interactions within the CHI3L1 binding pocket. The 2D interaction diagram of **K284** in complex with CHI3L1 predicted the formation of hydrogen bonds with Asp207, Tyr141 and arg263. Additional π-π stacking interactions and hydrophobic contacts with multiple residues of the binding site were observed. However, despite the otherwise extensive network of favorable interactions, **K284** demonstrated an unfavorable interaction (Figure 6E) with Glu290 amino acid residue of CHI3L1. The presence of an unfavorable interaction with Glu290 points to a potential limitation in the binding efficacy of **K284** and helps to explain to the observed modest efficacy in the MST assay.

Molecular docking of the reported compound **G721** (depicted in red, Figures 6F-G) showed that the compound established several π-π stacking interactions and a π-sulfur interaction with the amino acid residues of the CHI3L1 binding site. However, G721 did not form any hydrogen bonds with the binding site residues, highlighting the potential importance of hydrogen bonding for stable and potent binding. Lastly, **G28** (illustrated in blue, Figures 6H-J) exhibited the most extensive interaction network among the three investigated compounds correlating with observed results from the MST assay. The 2D diagram revealed the formation of multiple hydrogen bonds with residues Tyr141 and His209 amino acid residues of the binding site. Additionally, **G28** forms π-π stacking interactions as well as π-sulfur interactions indicating a diverse binding mode.

Overall, the molecular docking investigation reveals that the presence of hydrogen bonds and an extensive network of favorable interactions, coupled with the absence of unfavorable interactions, are critical determinants for identifying potent CHI3L1 binders. Additionally, the docking results convincingly demonstrate that molecular docking is a robust and reliable technique for screening and filtering potential CHI3L1 inhibitors, as evidenced by the distinct interaction profiles captured for **CHI3L1-IN-1**, **K284**, **G721** and **G28**. This success further validates the computational approach chosen for this project, providing a solid foundation for the rational design of more potent and selective CHI3L1 inhibitors with optimized pharmacological properties. The high spatial complementarity observed between the investigated compounds and the binding pocket reinforces the utility of structure-based virtual screening as an efficient strategy for identifying novel therapeutic candidates targeting CHI3L1-mediated pathological processes.

Following the docking studies, 100 molecular dynamics (MD) simulations were carried out to validate the predicted binding mode of the best hit, compound **G28** in comparison to the reported compounds **CHI3L1-IN-1** and **K284** which demonstrated dose-dependent binding. Additionally, a separate MD simulation of the unbound CHI3L1 protein served as a control. The structural stability of the investigated protein/ligand complexes in comparison to the unbound CHI3L1 protein was assessed by plotting the root-mean-square deviation (RMSD) over the 100ns duration of the MD simulations (Figure 7A). The unbound CHI3L1 exhibited substantial fluctuations over the duration of the 100ns MD simulation with RMSD values rising beyond 6 Å which indicates significant conformational flexibility. In contrast, all ligand-bound CHI3L1 complexes displayed a markedly lower and more stable RMSD values with average RMSD values of 1–2 Å. All three investigated complexes, **CHI3L1-IN-1**/CHI3L1, **G28**/CHI3L1 and **K284**/CHI3L1 complexes demonstrate structural stability with low RMSD values which suggest that ligand binding significantly enhances the conformational stability of CHI3L1.

**Figure 7.**
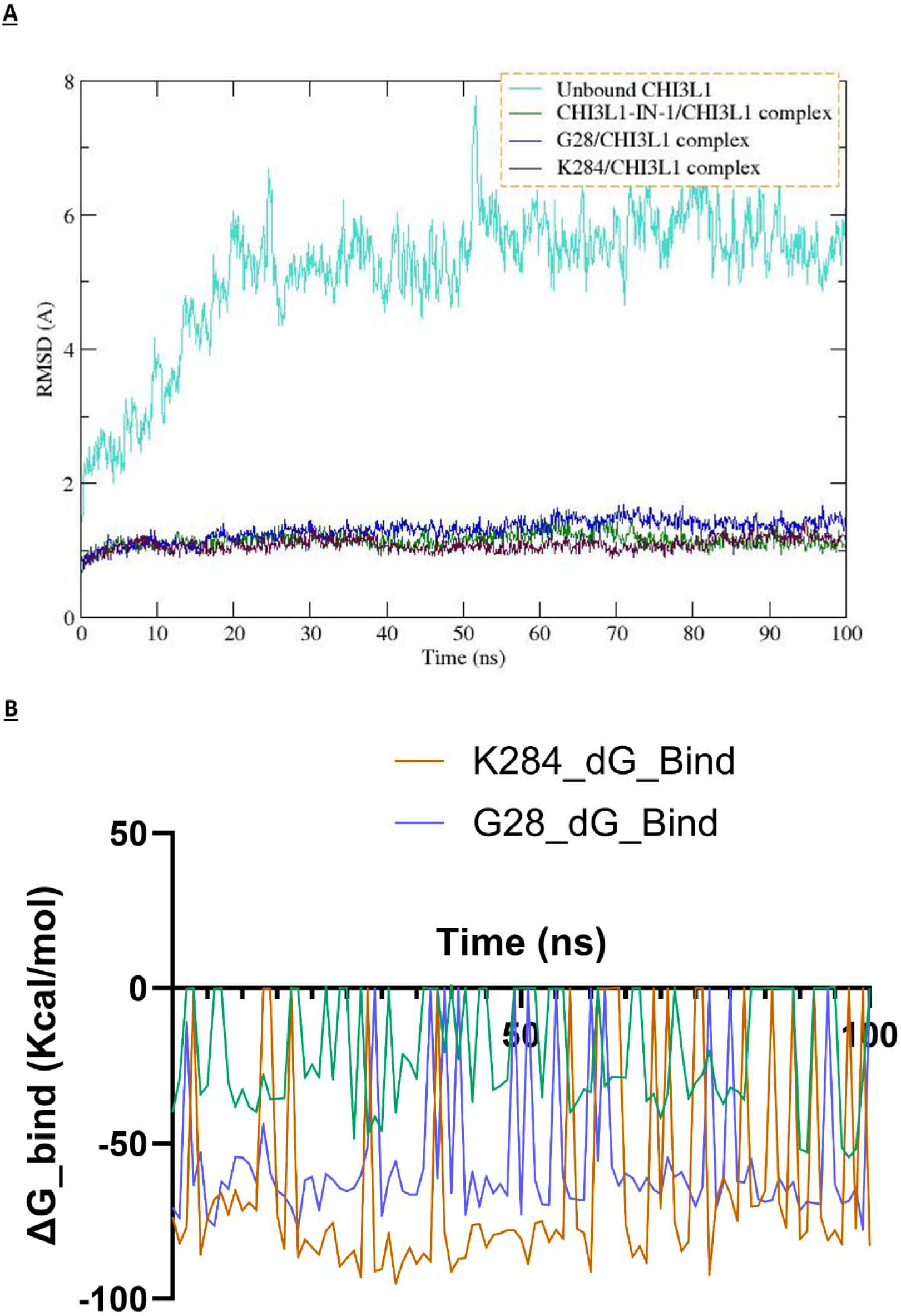
MD analysis of CHI3L1 in complex with CHI3L1-IN-1, K284 and **G28.** (A) RMSD of CHI3L1 in complex with **CHI3L1-IN-1**, **K284**, **G28** over 100 ns, showing system equilibration and structural stability with fluctuations centered around ∼1.2 Å. (B) Free energy analysis of the CHI3L1 in complex with **CHI3L1- IN-1**, **K284 and G28** across the duration of the 100ns MD simulations.

Interaction analysis (Figure S4) demonstrates that all three investigated compounds maintained consistent contact with Trp99 throughout the 100 ns MD simulations, indicating a shared binding mode and establishing Trp99 as a critical residue for ligand recognition and stabilization. Analysis of binding free energy (ΔG_bind) across the CHI3L1 complexes revealed that **CHI3L1-IN-1** exhibited the highest (least favorable) average free energy at -20.4 kcal/mol. In contrast, **K284** and **G28** displayed substantially more favorable binding energies of -63.0 and -55.5 kcal/mol, respectively. These computational values correlate strongly with experimental binding data, where **K284** and **G28** demonstrated significantly higher potency than **CHI3L1-IN-1**. Moreover, plotting the free energy analysis for the investigated compounds over the duration of the MD simulations (Figure 7b) showed that **CHI3L1-in-1** possessed highly fluctuating free energy values which is an indicator of transient instability. Conversely, both **K284** and **G28** demonstrated a significantly stronger and more stable binding profile, although a few frames reveal anomalously high (less negative or even near-zero) values which is a possible indicator of transient binding. Together, these results illuminate the binding mechanisms of the investigated compounds and further demonstrate that computational techniques can effectively discriminate between active and inactive CHI3L1 binders.

### 2.5. PK profiling

The development of systemically administered small molecule therapeutics for GBM relies not only on target specificity and potency but also on optimal pharmacokinetic (PK) properties that ensure sufficient exposure in the brain and tumor microenvironment. In this context, we performed a comparative in vitro ADME evaluation of four compounds (**G28**, **CHI3L1-IN-1**, **K284**, and **G721-0282**). The in vitro PK evaluation of **G28** and previously reported CHI3L1 small molecule inhibitors (**CHI3L1-IN-1**, **K284**, and **G721-0282**) highlights the markedly favorable profile of **G28** across key ADME parameters (Tables 1 and 2). **G28** demonstrated the longest half-life (Table 1) in both mouse and human plasma (3.1 h and 3.9 h, respectively), indicating enhanced stability in systemic circulation compared to **CHI3L1-IN-1** (0.9–1.2 h), **K284** (1.5–1.9 h), and **G721-0282** (0.7–0.8 h). This increased plasma stability is critical for achieving sustained drug levels in vivo and reducing the frequency of dosing, an especially important consideration for GBM patients who often experience rapid disease progression and may have limited tolerance to frequent drug administration. **G28** also displayed markedly improved microsomal stability (Table 1), with half-lives of 3.6 h (mouse) and 4.0 h (human), translating into low intrinsic clearance values (12 and 8.1 mL/min/mg, respectively). These results suggest reduced hepatic metabolic liability, which in turn supports a favorable systemic exposure profile. In contrast, **CHI3L1-IN-1**, **K284**, and **G721- 0282** demonstrated higher clearance rates, indicating a higher likelihood of rapid metabolism and poor bioavailability in vivo.

**Table 1.** Assessment of the metabolic/microsomal stability of **G28, CHI3L1-IN-1, K284**, and **G721-0282**.

| Compound | $t_{1/2}$ [h]<br>(mouse<br>plasma) | $t_{1/2}$ [h]<br>(human<br>plasma) | $t_{1/2}$ [h]<br>(mouse<br>microsomes) | $t_{1/2}$ [h]<br>(human<br>microsomes) | $Cl_{int}$ (mouse,<br>mL/min/mg) | $Cl_{int}$ (human,<br>mL/min/mg) |
| --- | --- | --- | --- | --- | --- | --- |
| **G28** | 3.1 ± 0.3 | 3.9 ± 0.4 | 3.6 ± 0.2 | 4.0 ± 0.1 | 12 ± 1.4 | 8.1 ± 0.6 |
| **CHI3L1-IN-1** | 0.9 ± 0.1 | 1.2 ± 0.2 | 1.1 ± 0.2 | 1.5 ± 0.4 | 28 ± 3.8 | 32 ± 4.2 |
| **K284** | 1.5 ± 0.2 | 1.9 ± 0.3 | 1.7 ± 0.4 | 2.1 ± 0.3 | 18 ± 1.6 | 22 ± 1.3 |
| **G721-0282** | 0.7 ± 0.1 | 0.8 ± 0.1 | 0.8 ± 0.2 | 0.9 ± 0.1 | 42 ± 5.3 | 45 ± 2.7 |

<sup>a</sup>Values are mean ± SD (n=3).

**Table 2.** Assessment of key ADME properties of **G28, CHI3L1-IN-1, K284**, and **G721-0282**.

| Compound | Kinetic Solubility 1% DMSO/PBS [$\mu M$]<sup>a</sup> | PAMPA permeability (cm/s) | LogD<sub>7.4</sub><sup>b</sup> | IC<sub>50</sub> against hERG [$\mu M$] | PPB human [%] |
| --- | --- | --- | --- | --- | --- |
| **G28** | 251 | $7.1 \times 10^{-6}$ | 1.8 | >100 | 79.5 |
| **CHI3L1-IN-1** | 53 | $2.2 \times 10^{-6}$ | 2.7 | $16 \pm 2.5$ | 96.1 |
| **K284** | 89 | $3.1 \times 10^{-6}$ | 2.1 | $22 \pm 3.1$ | 91.5 |
| **G721-0282** | 12 | $1.5 \times 10^{-6}$ | 3.3 | $5.3 \pm 0.8$ | 98.3 |

<sup>a</sup>Kinetic solubility measured in 1% DMSO/PBS (pH 7.4) by UV-vis spectrophotometry (n=3). <sup>b</sup>LogD (pH = 7.4) the distribution coefficient at physiological pH 7.4.

Beyond metabolic stability, **G28** possesses favorable drug-like properties (Table 2) that further support its prioritization for GBM therapy development. It exhibited the highest kinetic solubility (251 µM) in PBS/DMSO among the tested compounds, a critical factor for formulation and oral absorption. Moreover, **G28** showed the highest passive permeability in the PAMPA assay (7.1 × 10⁻⁶ cm/s), suggesting a strong potential for BBB penetration—an essential characteristic for any drug candidate intended to treat intracranial tumors. **G28** also demonstrated an optimal lipophilicity profile (LogD₇.₄ = 1.8), which balances membrane permeability with aqueous solubility and may reduce the likelihood of off-target CNS effects or accumulation in fatty tissues. Importantly, **G28** showed no measurable inhibition of the hERG potassium channel (IC₅₀ >100 µM), minimizing the risk of cardiotoxicity—a common liability that limits the clinical viability of numerous small molecules. Notably, **G28** also revealed a lower extent of plasma protein binding (79.5%) compared to **CHI3L1-IN-1** (96.1 %), **K284** (91.5%), and **G721-0282** (98.3%), which may translate to a higher unbound fraction in plasma and enhanced tissue distribution, including potential for improved CNS exposure—an important consideration in the context of GBM therapy.

To assess the cytotoxic potential of the tested compounds and support their safety as candidates for GBM therapy, we evaluated cell viability in three non-cancerous human cell lines: primary human astrocytes (NHA), human brain microvascular endothelial cells (hBMECs), and HepG2 hepatocellular carcinoma cells as a surrogate for hepatocytes (Figure 8). All compounds were tested at 50 µM for 72 hours. **G28** exhibited no significant cytotoxicity across all three cell types, maintaining >90% cell viability, indicating a favorable safety profile. In contrast, **CHI3L1-IN-1** and **G721-0282** significantly reduced cell viability, particularly in astrocytes and HepG2 cells, suggesting potential off-target effects. **K284** demonstrated moderate cytotoxicity in NHA, and acceptable viability in hBMECs and hepatocytes, suggesting it may be relatively safer among the non-lead candidates (Figure 8).

**Figure 8.**
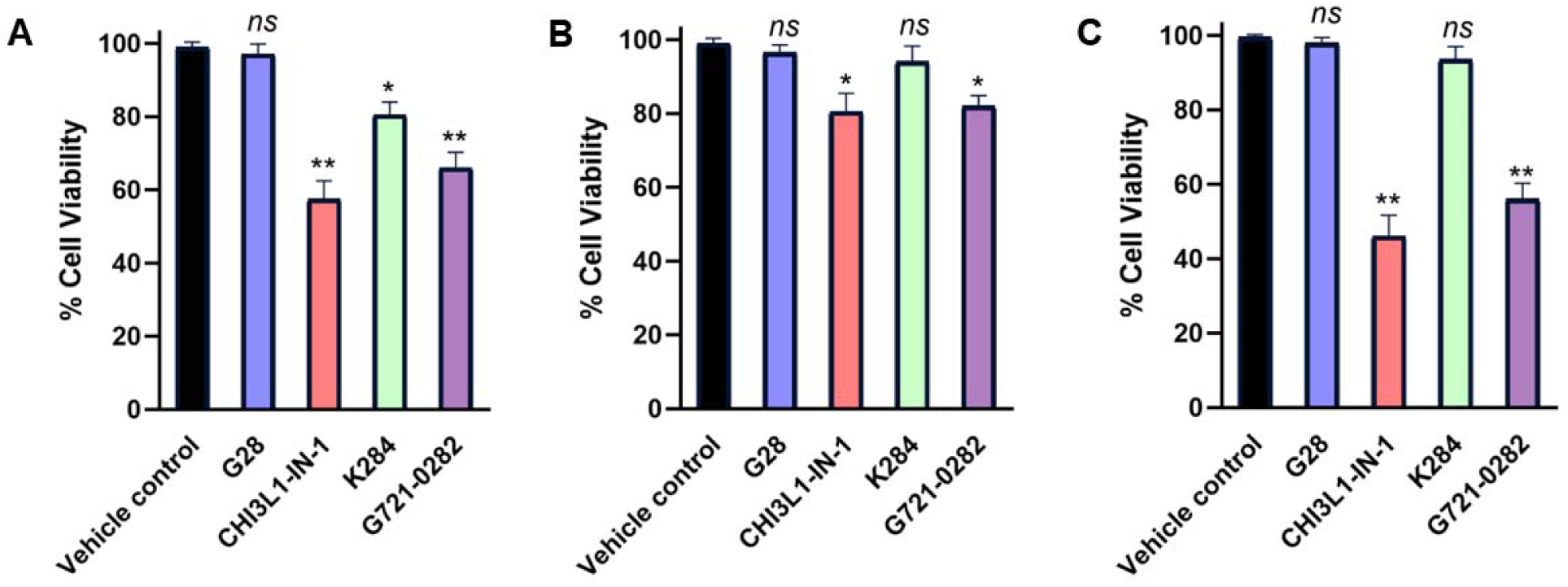
Cell viability as assessed by PrestoBlue of NHA (**A**), hBMECs (**B**), and HepG2 (**C**) upon incubation with a single dose of 50 μM of the tested compounds (**G28, CHI3L1-IN-1, K284**, and **G721-0282**) after 72 h incubation. * p < 0.05, ** p < 0.01, and (ns) denotes nonsignificant relative to vehicle control. Error bars represent standard deviation (n = 3).

Taken together, these data establish **G28** as a best-in-class compound as a small molecule CHI3L1 inhibitor, exhibiting the most favorable composite in vitro ADME profile. The enhanced profile of **G28** in comparison to the other investigated compounds becomes very apparent once the key ADM parameters are visualized via a radar plot (Figure 9). The spider plot configuration effectively illustrates how **G28** maintains an optimal combination of drug-like properties, with appropriate lipophilicity (LogD_₇.₄_ = 1.8) balancing solubility and membrane permeability, while exhibiting minimal hERG inhibition (IC₅₀ >100 µM) and lower plasma protein binding (79.5%) compared to the other investigated compounds. Overall, the enhanced stability, permeability, solubility, and safety margin exhibited by **G28** are highly advantageous for central nervous system (CNS) drug development and position it as a strong candidate for further in vivo pharmacokinetic and efficacy studies in GBM models. These PK properties, combined with its minimal toxicity support our conclusion that **G28** could overcome several of the barriers that have hindered small molecule-based CHI3L1 inhibition for GBM therapy.

**Figure 9.**
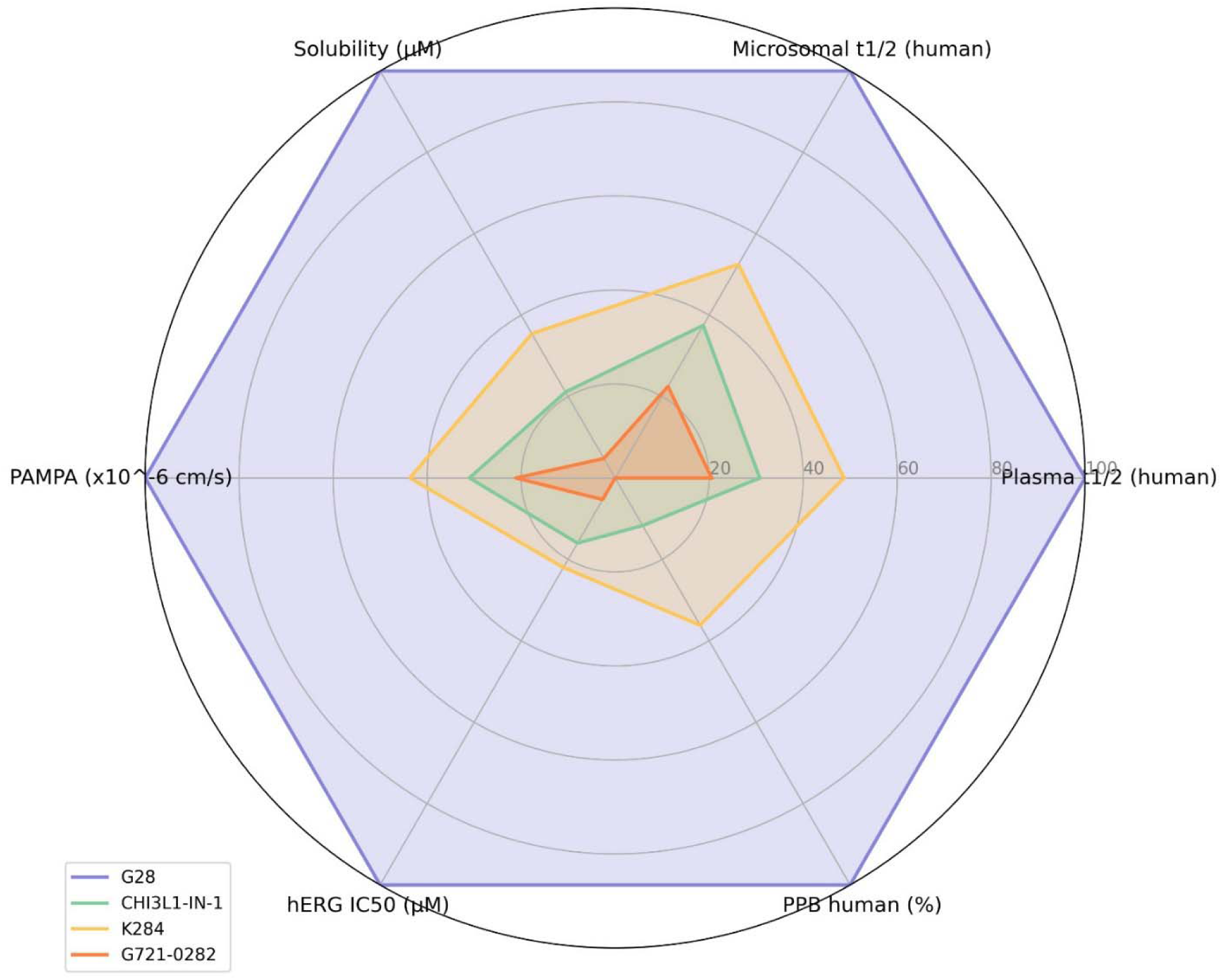
Comparative ADME profile of G28, CHI3L-IN-1, K284 and G721. Each parameter was normalized to a 0-100 scale. **G28** consistently outperforms the other investigated compounds across nearly all measured parameters.

### 2.6. Evaluation in GBM spheroids

Three-dimensional (3D) spheroid models of GBM have emerged as superior experimental platforms compared to traditional two-dimensional (2D) monolayer cultures. Unlike 2D cell cultures, which fail to replicate the complex architecture and cellular interactions of solid tumors, spheroid models better mimic the in vivo tumor microenvironment. This is particularly crucial for GBM, a highly aggressive and heterogeneous brain tumor characterized by a dense vascular network and significant immune cell infiltration. The spheroid structure allows for the formation of gradients in oxygen, nutrients, and signaling molecules, thereby more accurately simulating the physiological conditions of tumors. Additionally, the multicellular composition of spheroids enables the study of tumor-stroma interactions, which are critical for evaluating therapeutic responses.

We evaluated the efficacy of the CHI3L1-targeted compounds using a multicellular spheroid model comprising three key cell types typically found in brain tumors: glioblastoma cells, microvascular endothelial cells, and macrophages. This model is highly relevant for studying GBM, as these tumors are marked by extensive vascularization and significant macrophage infiltration—up to 40% of the tumor mass. These macrophages play a vital role in mediating the interplay between immune responses and angiogenesis within the tumor microenvironment. The reported GBM spheroid model used in our study^32^ is composed of U-87 MG glioblastoma cells, human microvascular endothelial cells (HMEC-1), and macrophages.

As shown in Figure 10, we first assessed compound efficacy by measuring cell viability in the GBM spheroids upon incubation with increasing concentrations of each tested compound. Among the tested compounds, **G28** exhibited the most pronounced reduction in spheroid viability in a dose-dependent manner, with statistically significant effects at lower concentrations compared to the other compounds (Figure 10). **K284** showed a modest, yet measurable, decrease in viability, whereas **CHI3L1- IN-1** and **G721-0282** had no significant impact, suggesting a lack of cytotoxicity or insufficient target engagement in this 3D context (Figure 10).

**Figure 10.**
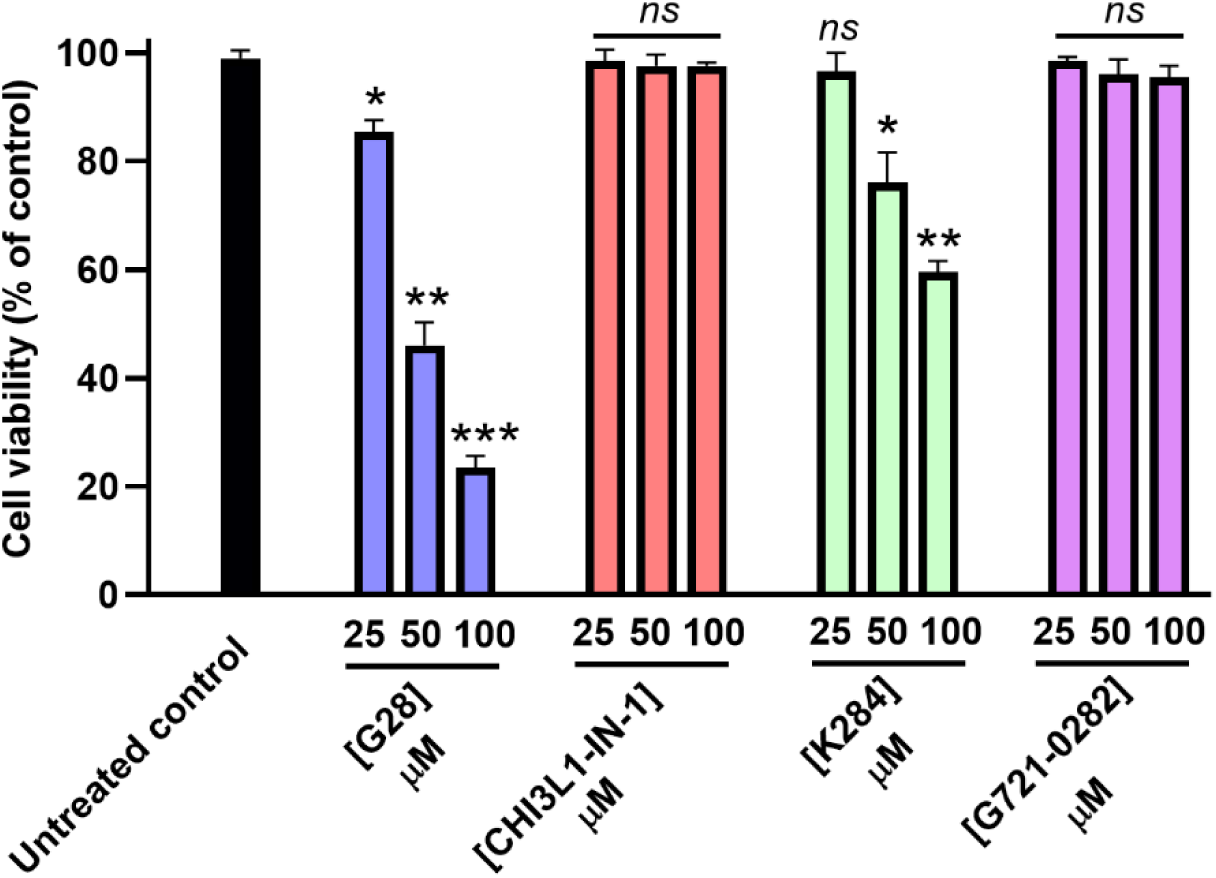
Cell viability of GBM spheroids (as % of untreated control) upon incubation with increasing concentrations of the tested compounds (**G28, CHI3L1-IN-1, K284**, and **G721-0282**) after 72 h incubation. * p < 0.05, ** p < 0.01, *** p < 0.001, and (ns) denotes nonsignificant relative to untreated control. Data are representative of three independent experiments.

To validate these findings, we measured the physical weight of spheroids after 72 hours of incubation with a single dose (50 μM) of each tested compound. **G28** again exhibited the most pronounced effect, inducing a significant reduction in spheroid mass, which is consistent with reduced cellular density and may reflect compromised structural integrity of the multicellular spheroid architecture (Figure 11). The weight reduction mirrored the viability trends, further supporting the cytotoxic activity of **G28**. **K284** induced a less pronounced, though statistically significant, decrease in spheroid weight, while the remaining two compounds (**CHI3L1-IN-1** and **G721-0282**) produced no significant changes, reinforcing their inactivity (Figure 11).

**Figure 11.**
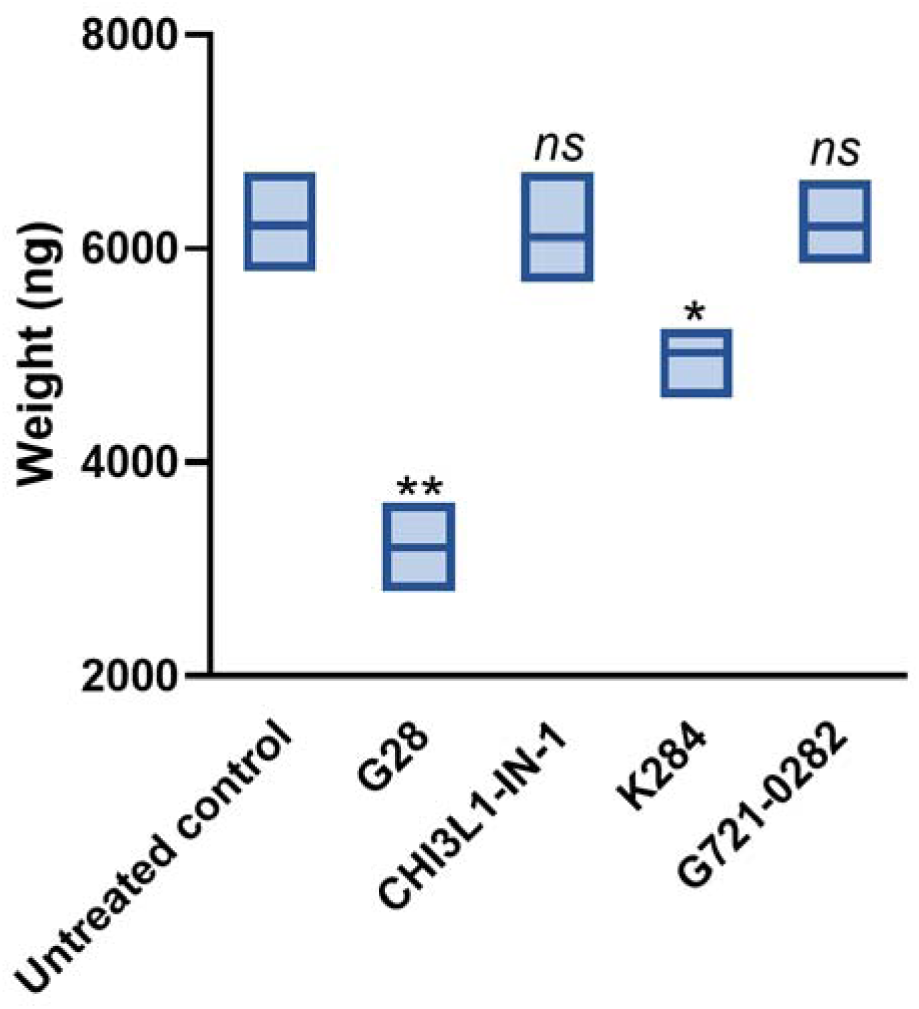
Changes in the weight of GBM spheroids upon incubation with 50 μM of the tested compounds (**G28, CHI3L1-IN-1, K284**, and **G721-0282**) after 72 h incubation. * p < 0.05, ** p < 0.01, and (ns) denotes nonsignificant relative to untreated control. Data are representative of three independent experiments.

Finally, we evaluated the ability of compounds (at 50 μM) to inhibit spheroid outgrowth, serving as a surrogate measure of tumor invasiveness. As shown in Figure 12, **G28** significantly impaired migration from the spheroid core, indicating that in addition to its cytotoxic effects, it may interfere with GBM cell motility or microenvironmental signaling pathways that facilitate invasion. **K284** had a limited effect on migration distance, while **CHI3L1-IN-1** and **G721-0282** failed to alter migratory behavior (Figure 12). Taken together, these data position **G28** as the most promising CHI3L1-targeted candidate reported to date as a potential candidate for GBM therapy, with consistent efficacy across multiple functional endpoints. The inclusion of endothelial and immune cell types in the spheroid model provides a more physiologically relevant system and suggests that **G28** functions within the broader cellular network that sustains tumor progression.

**Figure 12.**
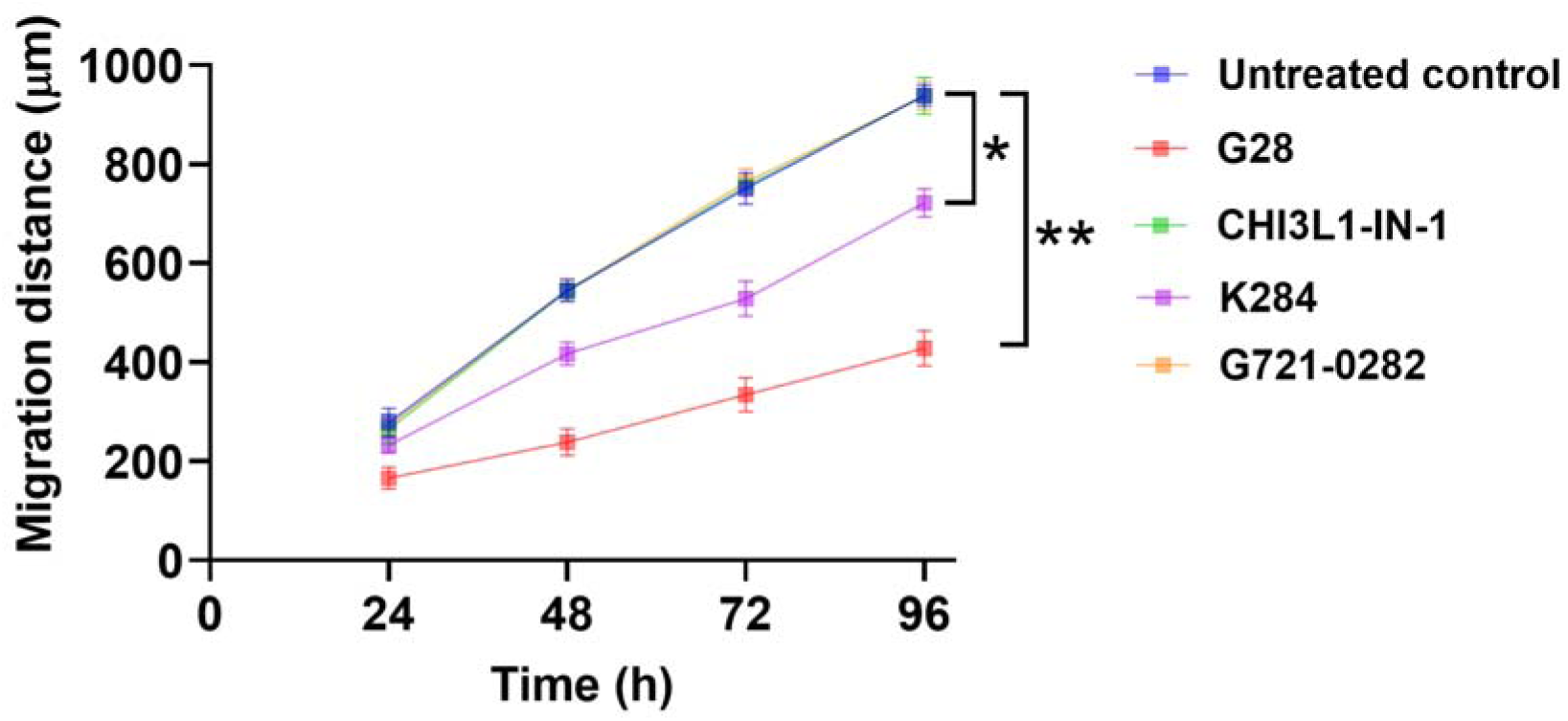
Cell migration distance (μm) of GBM spheroids upon incubation with 50 μM of the tested compounds (**G28**, **CHI3L1-IN-1**, **K284**, and **G721-0282**) after 96 h incubation. * p < 0.05, ** p < 0.01, and (ns) denotes nonsignificant relative to untreated control. Data are representative of three independent experiments.

## 3. Materials and methods

### 3.1. VS, molecular docking and MD

The computational studies against CHI3L1 were performed using the Schrödinger suite 2021.2 (Schrödinger, Inc., USA). **CHI3L1-IN-1** which was reported as the most potent CHI3L1 binder to date was selected for pharmacophore generation. Molecular dynamics (MD) simulations were conducted using DESMOND software. Results visualization was accomplished through Discovery Studio Visualizer, QtGrace, and Visual Molecular Dynamics (VMD) software. The crystal structure of CHI3L1 complexed with **CHI3L1-IN-1** was retrieved from the Protein Data Bank (PDB ID: 8R4X). Protein preparation was carried out using the protein preparation wizard in Maestro under the default conditions.

An in-house library comprising 16,922 compounds was processed using the Maestro ligand preparation module. The ligand preparation workflow involved importing 3D structures into Maestro and optimizing geometry using molecular mechanics (MM). Next, the compounds are adjusted to physiological pH (7.4) and subjected to geometry optimization which is followed by energy minimization using the LigPrep module with the OPLS3 force field. Pharmacophore generation was carried out by docking **CHI3L1-IN-1** into the CHI3L1 active site employing the extra precision module of Maestro. The docked complex was then used to develop a pharmacophore via the "Develop Pharmacophore from protein-ligand complex" wizard in Maestro. The generated hypothesis was validated by evaluating score differences between active compounds (14 compounds from the same experimental series as **CHI3L1-IN- 1**) and non-active compounds. Decoy compounds (50 in total) were generated using the DUD-E online service (http://dude.docking.org/) based on the CHI3L1-IN-1 structure. The pharmacophore-based screening and subsequent HTVS were carried out according to the previously reported methodology^43^.

MD simulations were performed using the DESMOND simulation package where the simulation system was constructed in an automatically defined orthorhombic box containing the solute molecule, solvated with explicit SPC water molecules. All simulations were conducted under physiological conditions, 300 K temperature and 1 bar pressure, with a total production run time of 100 ns and a positional relaxation time of 1 ps^44^. The OPLS 2005 force field^45^ was used for energy minimization of the solvated system. Electrostatic interactions were calculated using the Particle Mesh Ewald (PME) method, while non-bonded interactions were treated with periodic boundary conditions and a cutoff radius of 9.0 Å. System preparation involved a comprehensive six-step relaxation protocol. Subsequently, energy minimization, temperature, pressure and system equilibration were all done under the default settings of DEMSOND. Post-simulation trajectory analysis was carried out using XMGRACE^46^ (version 5.1) and GraphPad Prism to interpret structural and dynamic properties.

### 3.2. MST

#### 3.2.1. MST-based single-dosage screening

The Monolith His-Tag Labeling Kit RED-tris-NTA 2nd Generation (Cat. #MO-L018, NanoTemper Technologies, München, Germany) was used for the hCHI3L1-His protein (Cat. #CH1-H5228, Acro Biosystems, Newark, DE, USA) labelling according to the manufacturer’s instructions. Briefly, a 100 nM dye solution was mixed with 200 nM hCHI3L1-His in PBST buffer and incubated at room temperature (r.t.) in the dark for 30 minutes before each experiment.

The labeled hCHI3L1-His protein was then diluted in assay buffer and mixed with the test compound with final concentrations of 20 nM and 250 μM for the protein and compound, respectively. The mixture was incubated at r.t. for 30 minutes in the dark, followed by centrifugation at 1000 × g for 30 seconds before loading into the Dianthus NT.23 Pico instrument (NanoTemper Technologies, München, Germany). Assay buffer containing DMSO alone was used as the negative control. Assay buffer composition: 10 mM HEPES, 150 mM NaCl, 1% Pluronic F-127, 1 mM TCEP, 2.5% DMSO, pH 7.4. All assays were performed in technical triplicate, and results are reported as mean values.

#### 3.2.2. Autofluorescence test

Test compounds were diluted from DMSO stock solutions into assay buffer to a final concentration of 250 μM. The mixtures were incubated at r.t. for 30 minutes in the dark and centrifuged at 1000 × g for 30 seconds before loading into the Dianthus NT.23 Pico. Assay buffer containing DMSO alone served as the negative control. Measurements were performed in technical triplicate, and results are presented as mean ± standard deviation (SD).

#### 3.2.3. Quench test

Test compounds were mixed directly with RED-tris-NTA 2nd dye (Cat. #MO-L018, NanoTemper Technologies) at final concentrations of 250 μM (compound) and 10 nM (dye) in the assay buffer. The mixture was incubated at r.t. for 30 minutes in the dark, followed by centrifugation at 1000 × g for 30 seconds before loading into the Dianthus NT.23 Pico. Assay buffer containing 10 nM RED-tris-NTA 2nd dye and DMSO was used as the negative control. Measurements were conducted in technical triplicate, and data are reported as mean ± SD.

#### 3.2.4. Dose-dependent assay

Dose–response measurements of compounds identified as binders in the initial screen were conducted using the Monolith NT.115 instrument (NanoTemper Technologies, München, Germany). His-tagged CHI3L1 protein (Acro Biosystems, Cat. #CH1-H5228) was labeled with the RED-tris-NTA 2nd Generation His-Tag Labeling Kit (Cat. #MO-L018) in PBS with 0.05% Tween 20, following the manufacturer’s protocol. Compounds were serially diluted from 11mM to low nanomolar concentrations across 16 data points in assay buffer (101mM HEPES, 1501mM NaCl, 0.1% Pluronic F-127, 11mM TCEP, pH 7.4, and 8% DMSO). Final DMSO concentration was kept at 4% in the assay mixture. Following labeling, the protein was incubated with each compound at a 1:1 ratio for 30 minutes. Samples were then centrifuged at 10001×1g for 30 seconds prior to loading into standard capillaries. MST measurements were carried out using 60–80% LED excitation (red channel) and medium to high MST IR-laser power. Data were analyzed using MO.Affinity Analysis v2.3 software (NanoTemper Technologies).

For **K284**, the protein labeling was performed in 101mM HEPES, 1501mM NaCl, 0.1% Pluronic F- 127, pH 7.4 using the same RED-tris-NTA kit. Serial dilutions of **K284** were prepared in 101mM HEPES, 1501mM NaCl, 0.1% Pluronic F-127, 11mM TCEP, pH 7.4, with 8% DMSO to ensure a final DMSO concentration of 4% in the assay mixture. However, due to **K284**’s limited solubility in DMSO, saturation of binding could not be achieved. Increasing the concentration of K284 beyond a threshold led to precipitation, thus limiting the ability to generate a complete dose–response curve.

### 3.3. SPR screening

The interaction between **G28** and His-tagged CHI3L1 was determined by SPR (Biacore 8K, Cytiva). Kinetic measurements were run using a single cycle kinetic approach. Increasing concentrations of **G28** were injected over the prepared surface of the Sensor Chip after CHI3L1 immobilization. Experiments were conducted at 25°C. The results are presented as sensorgrams obtained after subtraction of the background response signal from a reference flow cell and from a control experiment with buffer injection. The obtained data were analyzed using BiacoreTM Insight Evaluation Software (Cytiva).

### 3.4. In vitro PK profiling

These studies were performed as we previously reported^47^. The procedures involve the determination of LogD7.4 values, microsomal stability, kinetic solubility, and cytotoxicity against a panel of cell lines. Solubility studies were performed using UV−vis spectrophotometry. PrestoBlue cell viability assay was used to assess the cell viability.

### 3.5. Screening in GBM spheroids

The GBM spheroids were prepared as previously described^32^. U-87 MG GBM cells (ATCC) were cultured in DMEM containing 4.5 g/L glucose and 2 mM L-glutamine, supplemented with streptomycin and penicillin. HMEC-1 (ATCC) were maintained in MCDB131 medium with 10 mM L-glutamine, 10 ng/ml FGF, and 1 µg/ml hydrocortisone. Birefly, U-87 MG and HMEC-1 cells were co-seeded with macrophages on low-adhesion 96-well plates at 2 × 10³ cells per well in 100 µl of the tested compounds at varying concentrations (25, 50, and 100 µM) or control media. After 72 hours, cell viability was assessed using the CCK-8 assay according to the manufacturer’s recommended protocol. Absorbance at 450 nm was measured using a microplate reader.

Migration assays were conducted using a serum-free flat surface migration method. Geltrex (Gibco) mixed with DMEM medium (1:50) was added to 12-well plates, incubated overnight at 36 °C, and the supernatant was removed. A single GBM spheroid was then seeded in each well and incubated for 90 minutes at 36 °C before adding fresh medium. Spheroids were pre-treated with 50 µM of the compound for 24 hours. Migration distances were monitored and recorded over three days using ImageJ, and cell death in migrating cells was assessed by Propidium Iodide staining after 72 hours. The W8 Physical Cytometer was employed to measure spheroid weight in a label-free and non-invasive manner. Data were generated from three independent experiments.

## 4. Conclusion

Virtual screening of our in-house library (16,922 compounds) identified four hits with CHI3L1 direct modulatory activity in a dose-dependent manner (**G28**, **E4**, **E11**, and **E15)**. Among the identified hits, **G28** demonstrated the best potency with a KD = 51.42 µM which was significantly higher than that of the previously reported modulators: **K284** (KD = 152 µM), **CHI3L1-IN-1** (KD = 77.3 mM), and **G721** (no dose-dependent binding detected). Computational analyses revealed strong correlations between predicted binding interactions, molecular dynamics stability and experimental CHI3L1 inhibition which validates our in-silico approach for targeted modulator identification. Comprehensive pharmacokinetic profiling showed that **G28** possesses exceptional drug-like properties which included optimal lipophilicity (LogD₇.₄ = 1.8), negligible cardiac liability (hERG IC₅₀ > 100 µM) and favorable plasma protein binding (79.5%) which was superior to that of the currently reported CHI3L1 modulators. In 3D GBM spheroid assays, **G28** exhibited dose-dependent suppression of tumor viability, mass, and invasive capacity, with efficacy substantially exceeding that of **K284** (moderate activity) and both **CHI3L1-IN-1** and **G721** which demonstrated minimal effect. These findings establish **G28** as the most promising CHI3L1-targeted small molecule identified to date, uniquely combining robust target engagement, favorable pharmacokinetics, and potent anti-GBM activity in GBM spheroids.

## Abbreviations Used

2D: (Two-Dimensional)
3D: (Three-Dimensional)
ADME: (Absorption, Distribution, Metabolism, Excretion)
BBB: (Blood-Brain Barrier)
CHI3L1: (Chitinase-3-Like Protein 1)
CNS: (Central Nervous System)
Fnorm: (Normalized Fluorescence)
GBM: (Glioblastoma)
GSC: (Glioma Stem Cell)
hBMEC: (Human Brain Microvascular Endothelial Cell)
HTVS: (High-Throughput Virtual Screening)
IL-13Rα2: (Interleukin-13 Receptor Alpha 2)
KD: (Dissociation Constant)
MD: (Molecular Dynamics)
MST: (Microscale Thermophoresis)
NHA: (Normal Human Astrocyte)
PBS: (Phosphate-Buffered Saline)
PDB: (Protein Data Bank)
PK: (Pharmacokinetic)
RMSD: (Root-Mean-Square Deviation)
SPR: (Surface Plasmon Resonance)
TME: (Tumor Microenvironment)
TRIC: (Temperature-Related Intensity Change)
VS: (Virtual Screening)

## CRediT authorship contribution statement

Hossam Nada: Writing – original draft, Formal analysis, Data curation, Conceptualization. Longfei Zhang: Data curation. Baljit Kaur: Data curation. Moustafa Gabr: Supervision, and Funding acquisition.

## Notes

The authors declare no competing financial interest.

## Acknowledgments

This work was supported by the National Institute of Neurological Disorders and Stroke under grant number R01NS136524 (PI: Gabr).

