## Supporting Information for "CHI3L1-Targeted Small Molecules as Glioblastoma Therapies: Virtual Screening-Based Discovery, Biophysical Validation, Pharmacokinetic Profiling, and Evaluation in Glioblastoma Spheroids"

AUTHOR ADDRESS

* To whom correspondence should be addressed:

**Table of content**

| Supplementary Figures | Pages 3-5 |
| --- | --- |
| Enamine libaray VS hits | Page 6 |
| Medchemexpress library VS hits | Page 7 |

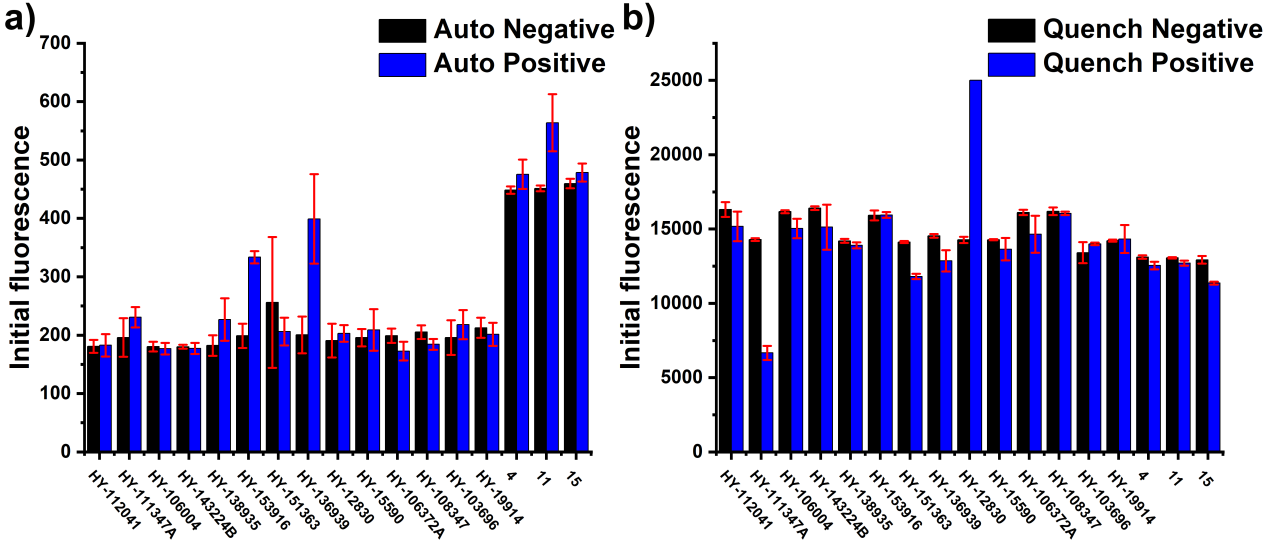

**Figure S1.** Autofluorescence (a) and quench test (b) of candidates from single-dosage screening.

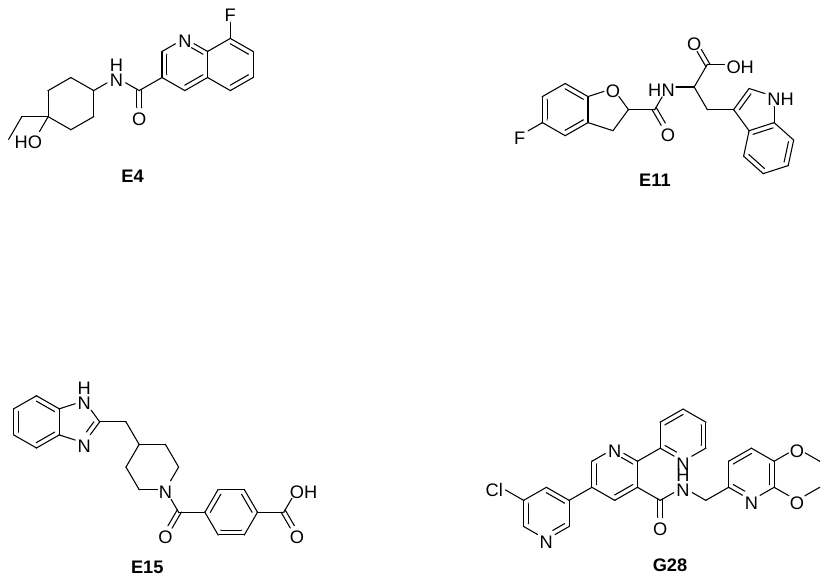

**Figure S2.** 2D structure of the top hit compounds from the VS study.

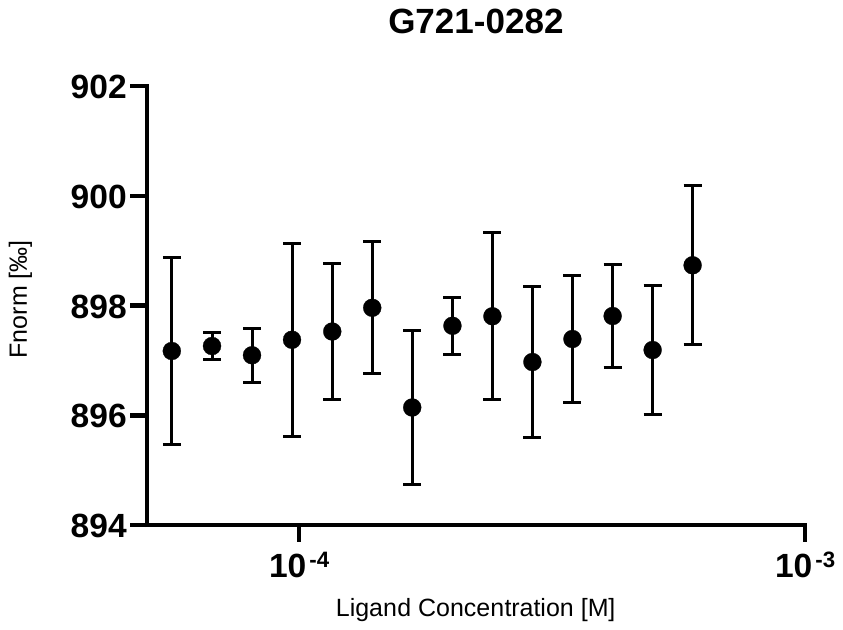

**Figure S3.** Direct binding evaluation of **G721** to CHI3L1 using MST.

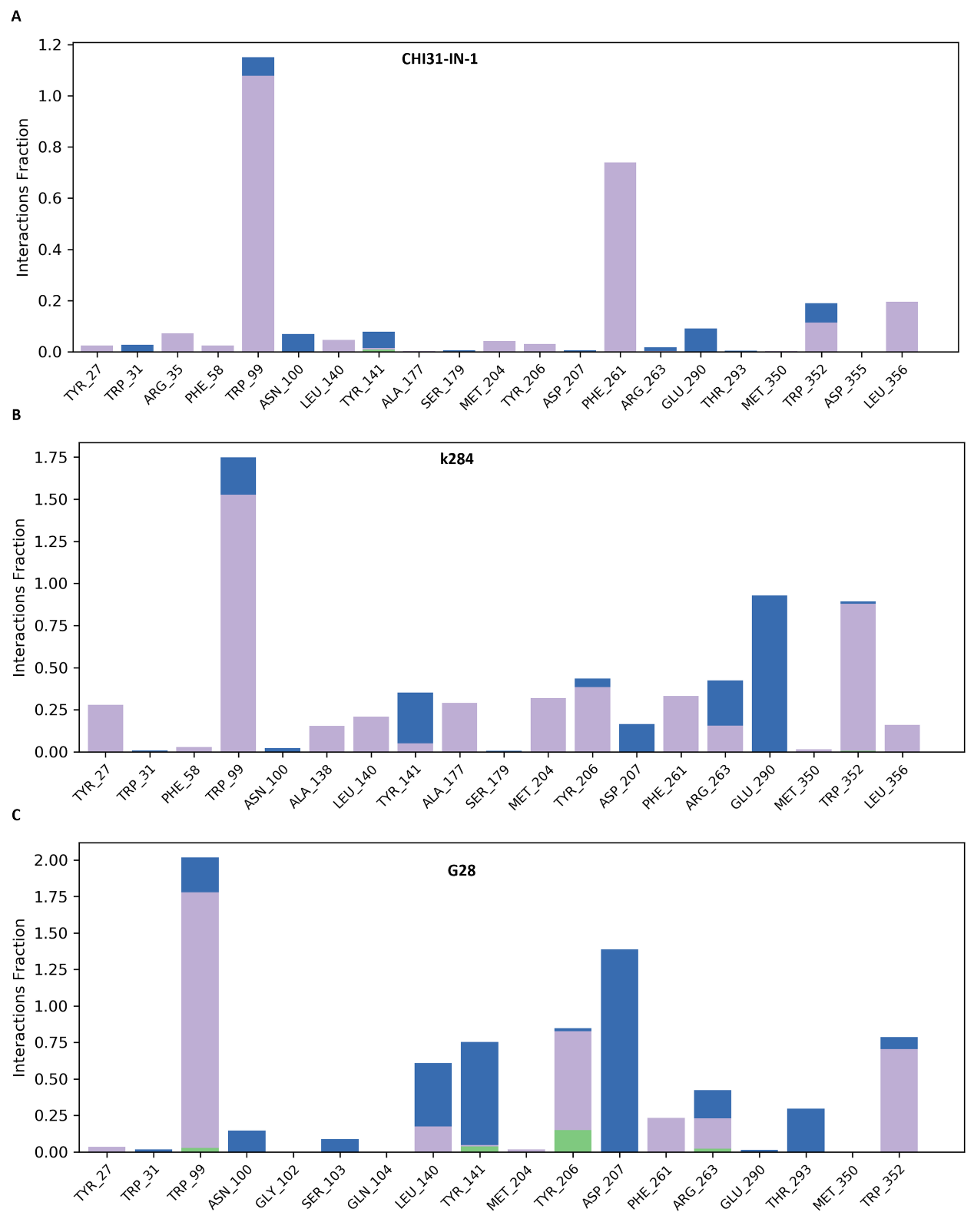

**Figure S3.** Protein-ligand contact histograms of CHI3L1 in complex with **CHI3L1-IN-1** (A), **K284** (b), **G28** (C).

**Table S1.** Enamine library VS hits subjected to VS.

|  | Catalog ID | SMILES | Manuscript ID |
| --- | --- | --- | --- |
| 1 | Z2771422053 | CC1=NN=C(S1)C(=O)N2CCC(CC2)NC(=O)C=3C=CC(C)=CC3 |  |
| 2 | Z1901266493 | CC1=NOC(CC(NC(=O)CC=2C=CC=C(C2)N3CCCC3=O)C=4C=CC=CC4)=N1 |  |
| 3 | Z752489918 | NC(=O)NCC1CCCN(C1)C(=O)NCC2CC3=CC=CC=C3O2 |  |
| 4 | Z2206759440 | CCC1(O)CCC(CC1)NC(=O)C2=CN=C3C(F)=CC=CC3=C2 | **E4** |
| 5 | Z1558922571 | CCC=1N=CN=C(C1F)N2CCN(CC2)C(=O)C=3C=CC=4NC=NC4C3 |  |
| 6 | Z1421305545 | CC=1C=C(C(=O)NCCOC=2C=CC(=CN2)C(F)(F)F)C=3C(=O)NN(C)C3N1 |  |
| 7 | Z2961428050 | CN1N=NN=C1C2CCN(CC2)C(=O)[C@@H]3C[C@H]3C=4C=CC(F)=CC4F \|&1:14,16,r\| |  |
| 8 | Z821876056 | COC=1C=CC=2NC=C(C2C1)C3CCN(CC4=NC(C)=C(C)O4)CC3 |  |
| 9 | Z804672914 | CC1=NOC(=N1)C2CCN(CC2)C(=O)CN3C=CC(=N3)C(F)(F)F |  |
| 10 | Z929239062 | O=C(O)COC=1C=CC=C(C1)NC(=O)C2CCN(CC2)C(=O)N3CCCC3 |  |
| 11 | Z1444487716 | O=C(O)C(CC1=CNC=2C=CC=CC12)NC(=O)C3CC4=CC(F)=CC=C4O3 | **E11** |
| 12 | Z1418887666 | NC(=O)C1CCC(CNC(=O)N2CCC(CC2)C3=NC4=CC=CC=C4S3)O1 |  |
| 13 | Z1468345650 | O=C(C=1C=CC(=CC1)N2C=NC=3C=CC=CC32)N4CCCC(C4)OCCO |  |
| 14 | Z2303756560 | COCCN1N=NC=2CN(CCC21)C(=O)C3=CN=C4C=CC=CC4=C3 |  |
| 15 | Z2442967527 | O=C(O)C=1C=CC(=CC1)C(=O)N2CCC(CC3=NC4=CC=CC=C4N3)CC2 | **E15** |
| 16 | Z2918016949 | CC1=NC=CN1CC(=O)N2CCC(CC3=NC(=NO3)C4=CC(C)=CC=N4)CC2 |  |
| 17 | Z2218794917 | CC(NC[C@@H](O)C1=CC=2C=CC=CC2O1)C=3C=CC=NC3 |  |
| 18 | Z1983369916 | CN1C(=NC=2C(F)=CC=CC21)N3CCC(CC3)C(=O)NCC=4C=CC(F)=CC4 |  |
| 19 | Z1619204556 | CC=1C=C(F)C=2N=C(O)C=C(C(=O)NC3CCCN(C=4C=CN(C)N4)C3=O)C2C1 |  |
| 20 | Z4015828039 | CCN1C=CC(CN(C)C(=O)C(=O)N[C@@H](C)C=2C=CC(=NC2)C(F)(F)F)=N1 |  |
| 21 | Z4076997412 | O=C(C1=CC(=NS1)C=2C=CC=CN2)N3CCCC(C3)C4=CC(O)=CC(F)=C4 |  |
| 22 | Z7077514455 | CC=1NN=CC1C(=O)NCC2CC(C)N(C2)C3=NC=4C=CC(F)=CC4O3 |  |
| 23 | Z1102431721 | CC1=NSC=2N=CC(=CC12)C(=O)N3CCN(CC3)C(=O)NC=4C=CC=C(C)C4 |  |
| 24 | Z1592804476 | CC(C)CN(CC(N)=O)C(=O)CCCC1=CC=C2C=CC=CC2=C1 |  |
| 25 | Z2384675768 | FC=1C=C(Cl)C=NC1NC[C@H]2CC[C@H](O2)C3=NC(=NN3)C4CC4 |  |
| 26 | Z3686109861 | O[C@@H]1CC(CN2C=C(Cl)C=N2)C[C@H]1NCC=3C=NC(=NC3)N4CCOCC4 \|&1:1,12,r\| |  |
| 27 | Z3687053243 | CC1=NN(C)C=2N=C(C=CC12)C(=O)N[C@@H]3CC(CN4C=C(Cl)C=N4)C[C@H]3O \|&1:14,25,r\| |  |
| 28 | Z3687120147 | NC(=O)C[C@@H]1CN(C[C@H]1O)C(=O)C=2C=CC=C(C2)CC=3C=CC=CN3 \|&1:4,8,r\| |  |
| 29 | Z5012236967 | OC=1C=C(C=CC1F)C2=NOC(=N2)C=3C=C(C=CN3)C=4C=NC=5NC=CC5C4 |  |

**Table S2.** Medchemexpress VS hits subjected to VS.

| Cat.# | 2D structure | Manuscript ID |
| --- | --- | --- |
| HY-112041 | NC1=C(F)C(NC2=CC=C(C(F)(F)F)C=C2)=NC(N3C4=CC(F)=CC=C4N=C3C)=N1 |  |
| HY-151096 | FC(C1=NC=C(C2=C(N3C[C@@H](C)N(C(CN4N=C(C)N=C4C)=O)CC3)SC(C(F)(F)F)=N2)C=N1)(F)F |  |
| HY-111347A | O=C(C1=CC=C(C2=CC=CC=C2)C=C1)N[C@H](C3=NC4=CC=CC=C4N3)CCCNC(CCl)=N.Cl |  |
| HY-114258 | O=C([C@@]1(CC2=NC(NC3=NNC(C)=C3)=CC=C2F)C[C@@H](C)N(CC4=CC=CC(Cl)=C4F)CC1)O |  |
| HY-106004 | S=C1N([C@@H]2CC3=CC(F)=CC(F)=C3OC2)C(CCNCC4=CC=CC=C4)=CN1 |  |
| HY-143224B | O=C(C(F)(F)F)O.CCC(C(NC1=CC=C(C=C1)F)=O)SC2=NC3=C(C4=NC(CCC5=C(NN=C5C)C)=NN24)C=CC=C3 |  |
| HY-138935 | O=C(C1=C(C=CC(S(C)(=O)=O)=C1)O[C@H](C)C(F)(F)F)N2C[C@]3(C4=NOC(C(F)(F)F)=C4)[C@@](C2)([H])C3 |  |
| HY-153916 | O=C(C1=CC(F)=CC=C1)NC2=CC=C(C3=C2C=CC=C3)OCC(NC4=CC=C(C(O)=C4)O)=O |  |
| HY-151363 | O=C([C@H]1[C@@H](C2)C=C[C@@H]2[C@H]1NC3=NC(NC4=CC=CC(C(N5CCCC5)=O)=C4)=NC=C3C6=CC(F)=CN=C6)N |  |
| HY-136939 | N#CC1=C(C2=CN=C(C(F)(F)F)N=C2)C3=C(N)N=CN=C3N1CC4=CN(C5=CC=CC=C5F)N=C4 |  |
| HY-153093 | CC(S1)=NN=C1CNC2=CC(OC[C@@H]3[C@@H](C4=CC=C(C=N4)C)C3)=NC(C)=N2 |  |
| HY-15651 | O=C(C1=CC(C2=CC=NN2C)=C(C)N(C3=CC=CC(C(F)(F)F)=C3)C1=O)NCC4=NC=C(S(=O)(C)=O)C=C4 |  |
| HY-15590 | CC(OC1=CC(NC2=NC(N[C@H](C3=NC=C(F)C=C3)C)=NC=C2Cl)=NN1)C |  |
| HY-114409 | O[C@H]([C@@H](N1N=NC(C2=CC(F)=C(F)C(F)=C2)=C1)[C@H]([C@@H](CO)O3)O)[C@H]3SC4=CC=C(Cl)C(Cl)=C4 |  |
| HY-10405 | O=C1C(OC2=CC=C(F)C=C2F)=CC3=CN=C(NC(CCO)CCO)N=C3N1C |  |
| HY-106372A | O=C(NC1=CC=CC=C1)C[N+](C)(C)CC(NC2=CC=CC=C2)=O.[Cl-] |  |
| HY-108347 | [H]Cl.COC1=CC2=NC(NCCC3=CC=C(OC)C(OC)=C3)=NC(N4CC5=C(C=C(OC)C(OC)=C5)CC4)=C2C=C1OC |  |
| HY-103696 | FC(C1=CC=C(NC2=NC(N3C4=CC(F)=C(F)C=C4N=C3C)=CN=C2)C=C1)(F)F |  |
| HY-19914 | O=C(C1=CC(C2=CC(Cl)=CN=C2)=CN=C1C3=NC=CC=C3)NCC4=NC(OC)=C(OC)C=C4 | **G28** |
